# Deacylation of SNAP25 protein family isoforms reveals distinct substrate selectivities of α/β hydrolase domain (ABHD) deacylases

**DOI:** 10.64898/2026.01.21.700842

**Authors:** Lorena Mejuto, Ludovico Pipito, Choa Ping Ng, Nicholas C.O. Tomkinson, Christopher A. Reynolds, Giuseppe Deganutti, Jennifer Greaves

## Abstract

Reversible *S*-acylation controls protein trafficking, localisation and activity within cells, yet the limited understanding of the regulatory mechanisms of S-acylation/deacylation cycles represents one of the biggest obstacles in exploiting this prevalent post-translational modification for therapeutic purposes. Here, we identified ABHD13 and ABHD16A as novel deacylases regulating the SNAP25 family of SNARE proteins responsible for vesicular membrane fusion. We determined the mechanism and structural determinants for ABHD13-mediated deacylation and substrate selectivity towards SNAP25b, and identified a novel mechanism of acyl chain extraction for integral membrane deacylases. This is the first report of deacylase selectivity mapped to a single amino acid position in full-length proteins.

## INTRODUCTION

Protein *S*-acylation (commonly ‘palmitoylation’), is the most prevalent and only reversible post-translational lipid modification, regulating thousands of cytosolic and transmembrane proteins^1–3^. Here, we identify the major brain phosphatidylserine (PS) lipase, ABHD16A^4^, and the mostly uncharacterised ABHD13, as thioesterases regulating the deacylation of the SNAP25 family of SNARE proteins, which are essential for vesicle fusion and exocytosis across neuronal and non-neuronal cells^5^.

*S*-acylation, the reversible addition of fatty acids onto cysteine residues, is critical for cell function, with dysregulated *S*-acylation-deacylation dynamics contributing to several cancers, metabolic and inflammatory diseases, as well as neurodegenerative and neuropsychiatric disorders through altered protein localisation, stability, and activity^2,6–8^. However, the promise of targeting *S*-acylation for therapy is limited by a fundamental lack of knowledge in how dynamic *S*-acylation is regulated by the enzymatic machinery.

*S*-acylation is reversed by *S*-deacylases that cleave the bond between the acyl chain and cysteine to remove the lipid.^9,10^ The majority of deacylases are metabolic serine hydrolases (mSH) and include the canonical deacylases Acyl Protein Thioesterase 1/2 (APT1/2), Palmitoyl Protein Thioesterase 1 (PPT1) and α/β-Hydrolase Domain 17 (ABHD17A/B/C)^10–16^. Additional mSH enzymes, including ABHD6, ABHD12, ABHD13 and ABHD16A, are also proposed to regulate dynamic *S*-acylation based on their sensitivity to deacylation inhibitors and activity-based probes^2,6,9,10,13,14^, but the exact number of *bona fide* deacylases is undetermined^2,6,14^.

*S*-acylation is particularly prevalent in the brain and nervous system, regulating almost half of all synaptic proteins^17^, including the neuronal SNARE (soluble *N*-ethylmaleimide-sensitive factor attachment protein receptor) protein SNAP25 (synaptic-associated protein of 25 kDa), where *S*-acylation is essential for stable membrane attachment and synaptic neurotransmitter release^18^. Moreover, SNAP25 *S*-acylation is altered in response to learning events in mice, and changes in the number of *S*-acylated sites control SNAP25 trafficking, subcellular localisation, association with cholesterol-rich membrane domains and exocytotic capacity, suggesting dynamic *S*-acylation is a regulatory mechanism for SNAP25 function^18–24^. While SNAP25 *S*-acyltransferases are known^23,25,26^, the deacylases involved have not been identified.

Despite the overall structural similarity within the mSH family, deacylases are highly selective for different *S*-acylated substrates and selective for distinct *S*-acylation sites within the same substrate^13,15,16,27–29^. The molecular basis for this selectivity is unknown, but likely involves substrate recognition by specific deacylases and their ability to access modified cysteines^11,15,27^. Following substrate recognition, the deacylases must identify the desired membrane-embedded *S*-acyl chain and then employ a mechanism to extract and sequester it. Recent short-timescale atomistic molecular dynamics simulations on APT2 and ABHD17, which are themselves *S*-acylated for stable membrane attachment, revealed that they insert a hydrophobic region into the membrane, perturbing the lipid bilayer and orienting a lipid binding pocket for optimal acyl chain entry^11,12^. However, the dynamic mechanism of acylated substrate recognition has not yet been demonstrated. Here, we present a structural model, informed by molecular dynamics (MD) simulations and validated by mutagenesis, of SNAP25 palmitoyl binding to ABHD13.

In summary, alongside identifying ABHD16A^4^ and ABHD13 as thioesterases that regulate SNAP25, we present evidence that ABHD13 has substrate specificity for different SNAP25 isoforms and distinct *S*-acylation sites, and that cysteine configuration determines favourable deacylation. Finally, we propose a novel mechanism for sequestering substrate acyl chains from the membrane. This association between substrate sequence and deacylase activity highlights a mechanism by which deacylases discriminate between individual *S*-acylated substrates and *S*-acylation sites.

## RESULTS

### ABHD13 and ABHD16A interact with SNAP25b to promote its deacylation and membrane displacement

Alternative splicing on exon 5 of the *Snap-25* gene results in two SNAP25 isoforms, SNAP25a and SNAP25b, which are differentially expressed during development^30^. SNAP25b, the main SNAP25 isoform expressed in human brains, is *S*-acylated at four cysteines within its membrane targeting domain^30,31^ (MTD; amino acids 85-120 - Fig. 1) by certain ZDHHC (Zinc finger Asp-His-His-Cys) *S*-acyltransferases^16,23,26,32^. SNAP25a is predominantly expressed during embryonic and postnatal development and differs from SNAP25b by nine amino acids (Fig. 1b)^30^. Whilst SNAP25a/b expression is limited to neuronal and neuroendocrine cells, the homologue, SNAP23, with 60% sequence identity, is ubiquitously expressed (Fig. 1b)^30^.

**Fig. 1:**
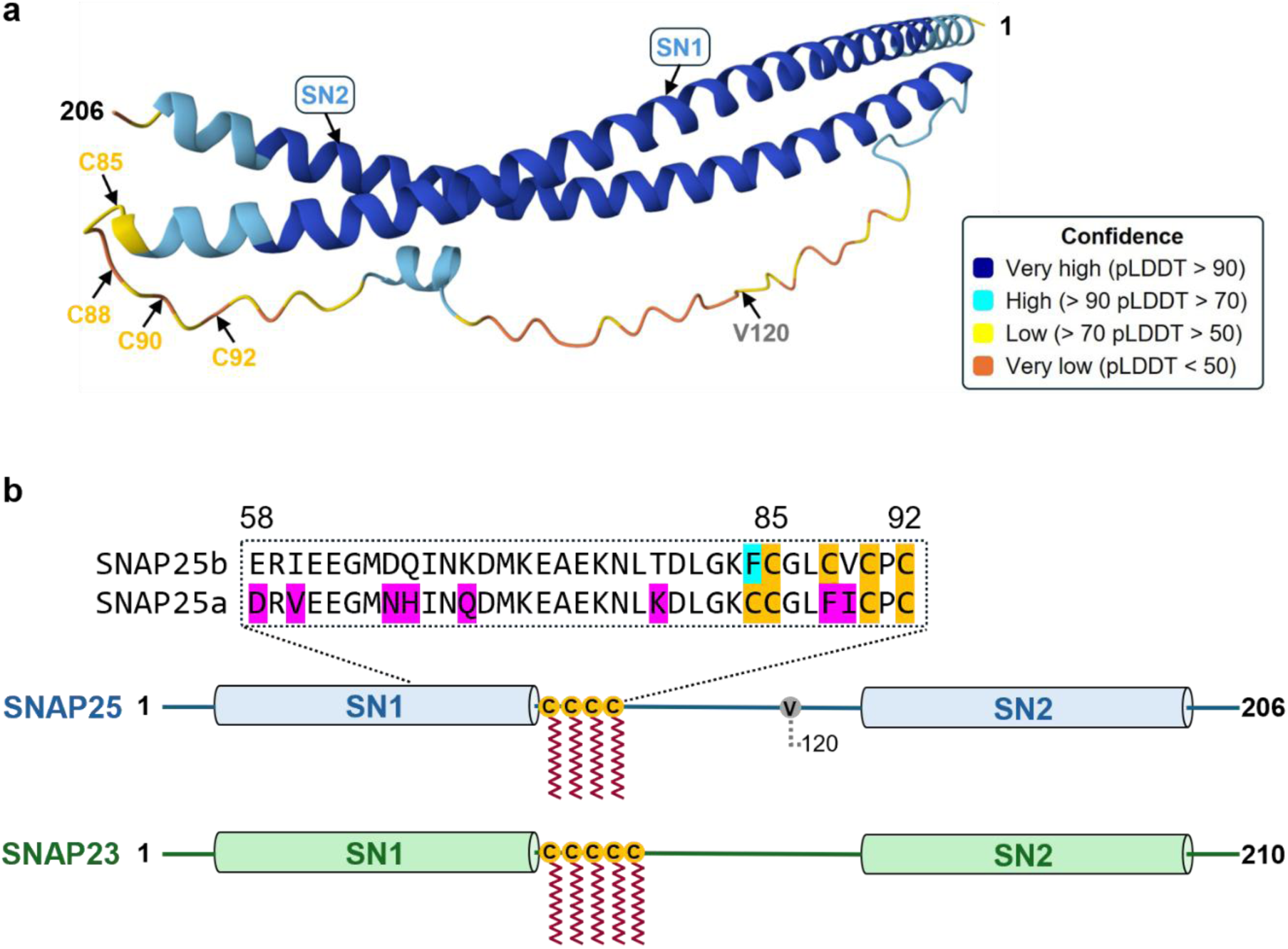
Schematic of the structure of the SNAP25 protein family. **a.** AlphaFold predicted structure of human SNAP25b (UniProt: P60880). Confidence (pLDDT score) is measured on a scale of 0-100. The two SNARE (SN) domains shown in blue are tethered together by the *S*-acylated flexible linker domain comprising four *S*-acylated cysteines (gold). The minimum membrane-binding domain comprises C85 through to V120^31^; the cysteine-rich domain (CRD) is C85 – C92. **b.** Cartoon diagram of human SNAP23 (green) and SNAP25a/b splice variants (blue). SNAP25 a/b splice variants differ by nine amino acids, highlighted in magenta and turquoise. V120, indicating the end of SNAP25b minimum membrane-binding domain, is shown in grey. SNAP25a/b contains up to four *S*-acylated cysteines, whereas SNAP23 contains up to five. *S*-acylated cysteines are highlighted in gold, and lipids are shown in red. A single phenylalanine (F84) is the only deviation in SNAP25b, not shared with SNAP25a or SNAP23 and is shaded in turquoise. The protein sequence alignment was generated using Clustal Omega^33^.

SNAP25b dynamic *S*-acylation is inhibited by the broad-range metabolic serine hydrolase (mSH) inhibitor Palmostatin B (PalmB), indicating that SNAP25b deacylation is enzymatically mediated by one or more PalmB-sensitive deacylases^13^; however, the enzyme(s) that deacylate SNAP25 proteins are unknown. HEK293T cells co-transfected with SNAP25b, ZDHHC3, APT1, APT2, ABHD10 or ABHD17A/B/C were metabolically labelled with the palmitic acid analogue C16:0-azide and conjugated to an IR dye by click chemistry to measure changes in SNAP25b *S*-acylation. We observed robust incorporation of the tagged palmitic acid into SNAP25b following co-transfection with ZDHHC3, but no observable change in *S*-acylation levels with APT1/2, ABHD10 or ABHD17A-C co-expression, suggesting that these deacylases do not target SNAP25b (Supplementary Fig. 2a-c). To investigate the role of other mSH enzymes in SNAP25b deacylation, we assessed SNAP25b *S*-acylation dynamics using metabolic labelling and pulse-chase analysis (Fig. 2a,b). HEK293T cells transfected with EGFP-SNAP25b and HA-ZDHHC3 were labelled with C16:0-azide for 4 hours, then chased with unlabelled palmitate for 3 hours in the presence of cycloheximide to block further protein synthesis. SNAP25b palmitate turnover was detectable in HEK293T cells and could be inhibited with PalmB, consistent with previous studies in COS-7 cells^13^. The ABHD6 inhibitor WWL70^34^ had no effect on SNAP25 deacylation, whereas tetrahydrolipstatin (THL), which additionally targets ABHD12 and ABHD16A^35^, prevented SNAP25b palmitate turnover to similar levels observed for PalmB (Fig. 2a,b), suggesting that ABHD12 or ABHD16A may be implicated in SNAP25b deacylation. Next, we assessed whether co-expression of ABHD6, ABHD13, or ABHD16A with EGFP-SNAP25b and HA-ZDHHC3 could alter SNAP25b *S*-acylation levels. ABHD13 has been shown to reduce postsynaptic scaffolding protein PSD-95 *S*-acylation levels moderately^16^. All three ABHD enzymes significantly reduced SNAP25b *S*-acylation, and there was no significant difference in the reduction of SNAP25b *S*-acylation between ABHD16A and ABHD13 (Fig. 2c-d). However, the exogenous expression of ABHD6 consistently lowered ZDHHC3 expression levels (Fig. 2c-e) and so was excluded from further analysis. The reduction in SNAP25b *S*-acylation by ABHD13 or ABHD16A was not due to alterations in ZDHHC3 activity, as *S*-acylation of SNAP25b by ZDHHC17 – a phylogenetically distinct SNAP25b *S*-acyltransferase to ZDHHC3^24,25^ – could also be reversed by ABHD13 or ABHD16A co-expression, indicating that the deacylase activity towards SNAP25b occurs independently of the initial SNAP25b *S*-acyltransferase (Supplementary Fig. 2d,e). Furthermore, these deacylases are substrate selective, since co-expression of ABHD16A or ABHD13 with the GTPase H-Ras, which is reversibly *S*-acylated on two cysteines^36^, had no detectable effect on its *S*-acylation levels (Supplementary Fig. 2f,g). Taken together, these results indicate that ABHD13 and ABHD16A are substrate-selective thioesterases that deacylate SNAP25.

**Fig. 2:**
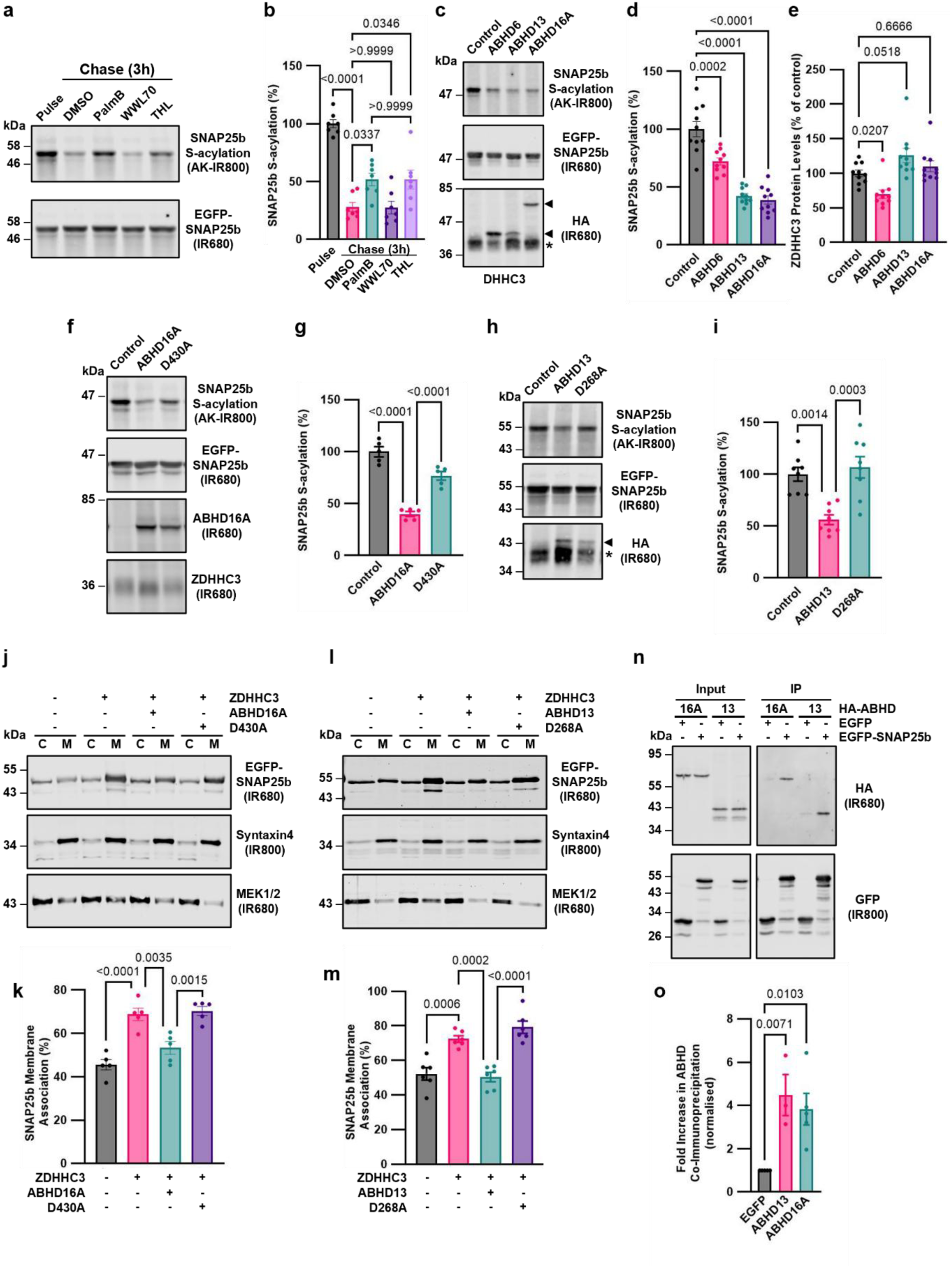
ABHD13 and ABHD16A interact with SNAP25b to promote its deacylation and membrane displacement. **a.** Pulse-chase analysis of C16:0-azide-labelled SNAP25b *S*-acylation turnover mock-treated with DMSO (control), or 100 μM PalmB, 10 μM WWL70 or 100 nM THL for 3 hours (3 h) analysed by click chemistry and immunoblotting. **b.** Quantification of SNAP25b *S*-acylation turnover in **a. c.** *S*-acylation levels of SNAP25b co-transfected with ZDHHC3 and empty vector (control), ABHD6, ABHD13 or ABHD16A determined by C16:0-azide-labelling, click chemistry and immunoblotting. **d.** Quantification of SNAP25b *S*-acylation levels in **c. e.** Quantification of ZDHHC3 protein levels in **c. f.** *S*-acylation levels of SNAP25b co-transfected with ZDHHC3 and empty vector (control), ABHD16A WT or D430A determined by C16:0-azide-labelling, click chemistry and immunoblotting. **g.** Quantification of SNAP25b *S*-acylation levels in **f. h.** *S*-acylation levels of SNAP25b co-transfected with ZDHHC3 and empty vector (control), ABHD13 WT or D268A determined by C16:0-azide-labelling, click chemistry and immunoblotting. **i.** Quantification of SNAP25b *S*-acylation levels in **h. j.** Membrane partitioning of HEK293T cells transfected with SNAP25b alone or with ZDHHC3 and empty vector or ABHD16A WT or D430A. **k.** Quantification of SNAP25b percentage membrane binding shown in **j. l.** Membrane partitioning of HEK293T cells transfected with SNAP25b alone or with ZDHHC3 and empty vector or ABHD13 WT or D268A. **m.** Quantification of SNAP25b percentage membrane binding shown in **l. n.** Co-immunoprecipitation of ABHD13 and ABHD16A from GFP-Trap® pull-down on SNAP25b. **o.** Quantification of mean fold increase in co-immunoprecipitated ABHD16A and ABHD13 compared to empty GFP vector shown in **n.**

We next identified amino acids critical for the deacylating activities of ABHD13 and ABHD16A. Using AlphaFold^37^ structural modelling we predicted the catalytic triad of ABHD13 and ABHD16A to comprise S193–H298–D268 and S355–H543–D430, respectively (Supplementary Fig. 2h; <u>U</u>niprot annotation for ABHD16A incorrectly named H507). To confirm this prediction and identify catalytically inactive enzymes, we created ABHD13(D268A) and ABHD16A(D430A) mutants. Binding of the fluorophosphonate (FP) probe azido-FP, which specifically and covalently labels the active-site serine of enzymatically active SHs, could be readily detected in WT ABHD16A; however, binding to the D430A mutant was completely abolished, indicating a catalytically dead enzyme (Supplementary Fig. 2i,j). Next, we determined whether the catalytic triad of ABHD13 and ABHD16A is required for SNAP25b deacylation. Co-expression of SNAP25b and ZDHHC3 alongside wild-type ABHD13, ABHD16A, or their catalytically inactive counterparts, combined with click chemistry analysis, showed that ABHD13 or ABHD16A deacylase activity towards SNAP25b requires an intact catalytic domain (Fig. 2f–i).

Previous work determined that SNAP25b membrane binding is inefficient in HEK293T cells but can be rescued by co-expression of certain ZDHHC *S*-acyltransferases^24^. However, it is unclear whether ABHD13- and ABHD16A-mediated deacylation of SNAP25b could reverse the effect of the *S*-acyltransferases in promoting membrane association. HEK293T cells expressing SNAP25b alone, with its *S*-acyltransferase ZDHHC3, or with ZDHHC3 and wild-type or catalytically dead deacylases (ABHD16A or ABHD13) were separated into cytosolic and membrane fractions to monitor SNAP25b membrane partitioning. Indeed, co-expression of both ABHD13 and ABHD16A, but not their catalytically dead mutants, was effective at reversing the effects of ZDHHC3-mediated SNAP25b membrane association (Fig. 2j-m and Supplementary Fig. 2. k,l). Furthermore, the interaction between ABHD13 and ABHD16A with SNAP25b could be detected by co-immunoprecipitation (Fig. 2n,o). Collectively, these results reveal that catalytically active ABHD13 and ABHD16A bind SNAP25b, catalysing its deacylation and membrane displacement in HEK293T cells.

### ABHD16A has greater activity towards SNAP25b than ABHD13

We next identified differences in ABHD13 and ABHD16A activity by examining dose-dependent effects on SNAP25b deacylation. We co-transfected HEK293T cells with SNAP25b, ZDHHC3 and increasing amounts (0 - 400 ng) of ABHD13 (Fig. 3a) or ABHD16A (Fig. 3b), followed by metabolic labelling and click chemistry analysis as before. The expression levels of ABHD13 and ABHD16A were quantified by dividing their signal intensity by that of ZDHHC3 at each data point and normalising these values to a maximum intensity of 100%. We plotted these values against the ABHD cDNA transfection concentration to generate a standard curve (Fig. 3c). Linear regression yielded coefficients of determination (R^2^) of 0.9346 for ABHD13 and 0.9728 for ABHD16A, confirming a linear relationship between transfection concentration and protein expression. Using these data, we quantified SNAP25b *S*-acylation levels at each concentration and plotted them relative to the control (100%). A non-linear dose-response curve showed that ABHD16A achieved a significantly higher level of SNAP25b deacylation at lower expression levels than that of ABHD13 (Fig. 3c,d). These results reveal notable differences in the activity of ABHD13 and ABHD16A toward SNAP25b deacylation under the same conditions.

**Fig. 3:**
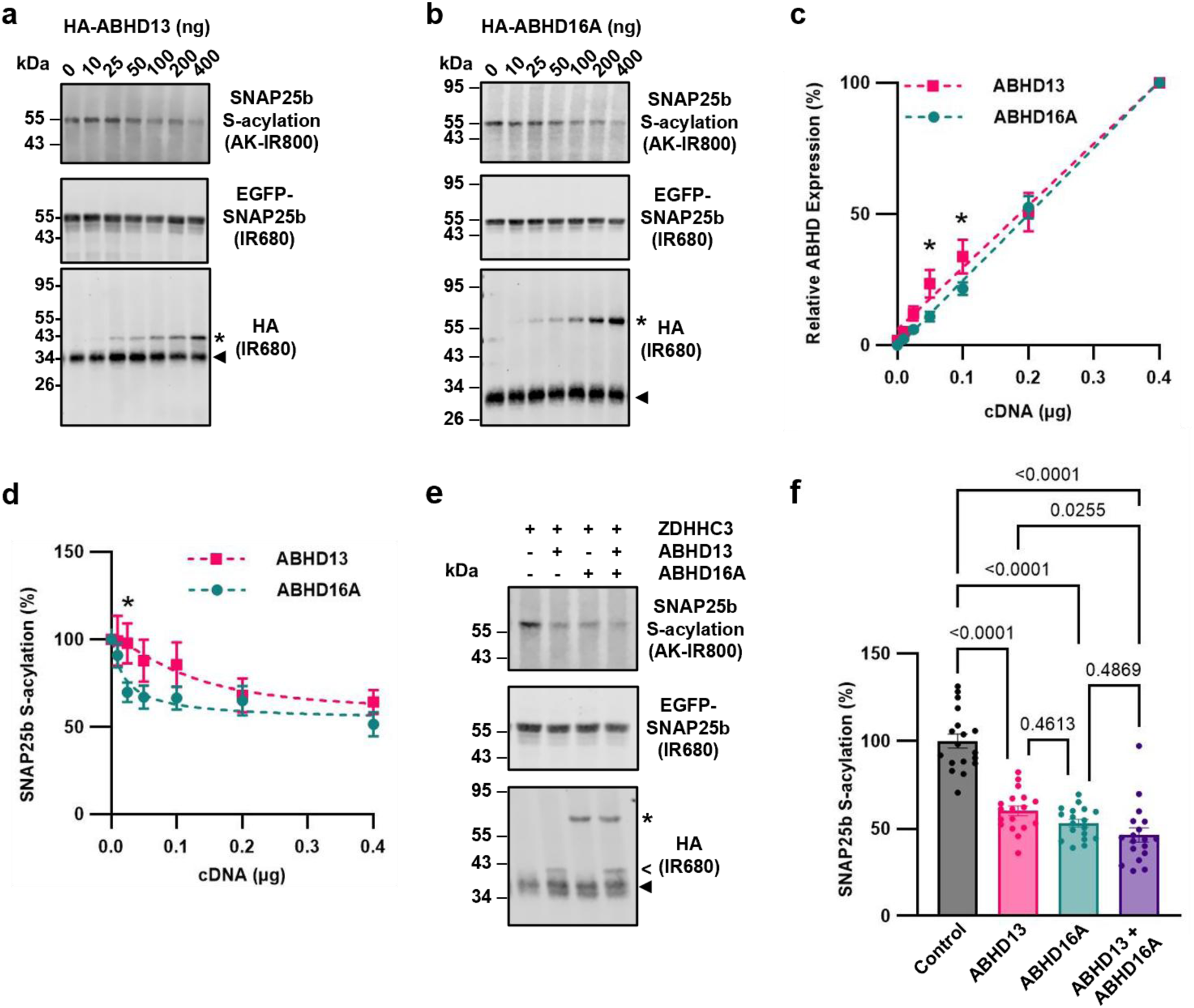
ABHD16A is more active than ABHD13 towards SNAP25b. **a, b.** Increasing amounts of co-transfected ABHD13 **(a.)** or ABHD16A **(b.)** plasmid reduce SNAP25b C16:0-az incorporation in a dose-dependent manner. **c.** Expression levels of ABHD13 (pink) and ABHD16A (teal) normalised to ZDHHC3 expression levels relative to the concentration of plasmid DNA transfected. **d.** Reduction in SNAP25b C16:0-az incorporation following co-expression of increasing concentration of ABHD13 (pink) or ABHD16A (teal), compared to control (empty vector). Quantification by unpaired t-test shown in **c.** and **d.**; *p < 0.05. **e.** *S*-acylation levels of SNAP25b co-transfected with ZDHHC3 and empty vector (control), ABHD13, ABHD16A or ABHD13 and ABHD16A determined by C16:0-azide-labelling, click chemistry and immunoblotting. **f.** Quantification of SNAP25b *S*-acylation levels in **e.**

Interestingly, neither enzyme could promote a loss in SNAP25b C16:0-azide labelling greater than ∼50%, even at the maximum expression levels tested (Fig. 3d). We explored whether the simultaneous expression of the enzymes – equivalent to a two-fold increase in exogenously expressed deacylase levels – might lead to further SNAP25b deacylation (Fig. 3e-f), but the reduction in C16:0-azide incorporation was only modest. These findings suggest that the SNAP25b pool reaches a steady-state equilibrium maintained by ZDHHC3 and ABHD activity.

### The SNAP25 isoforms SNAP25a and SNAP23 are displaced from membranes by ABHD16A-mediated deacylation only

Two studies by one group reported increases in immunoprecipitated SNAP23 from the supernatant of platelets following incubation with recombinantly purified APT1, but changes in the levels of SNAP23 *S*-acylation were not presented^38,39^. Differences in the configuration of *S*-acylated cysteines in the SNAP25 isoforms affect their interaction with the different ZDHHC *S*-acyltransferases^23^ and so we extended this study to investigate which exogenously expressed mSH could mediate SNAP25a and SNAP23 deacylation (Fig. 4a-d and Supplementary Fig. 3a,b). Notably, only the exogenous expression of ABHD16A could significantly reduce the *S*-acylation levels of SNAP25a and SNAP23 (Fig. 4a-d). Moreover, fractionation assays were used to quantify the degree of membrane association of SNAP25a or SNAP23 in the presence of ZDHHC3 and ABHD13, ABHD16A, or empty vector. This revealed that whilst ABHD16A-mediated deacylation reversed the effects of ZDHHC3 in promoting SNAP25a and SNAP23 membrane affinity, similar to SNAP25b (Fig. 2j-m), ABHD13 was not sufficient to displace SNAP25a and SNAP23 from membranes (Fig. 4e-h and Supplementary Fig. 3c,d). These results indicate that ABHD16A regulates the deacylation and membrane dissociation of all three SNAP25 isoforms, whereas ABHD13 can distinguish among the SNAP25 family members and prefers to deacylate SNAP25b.

**Fig. 4:**
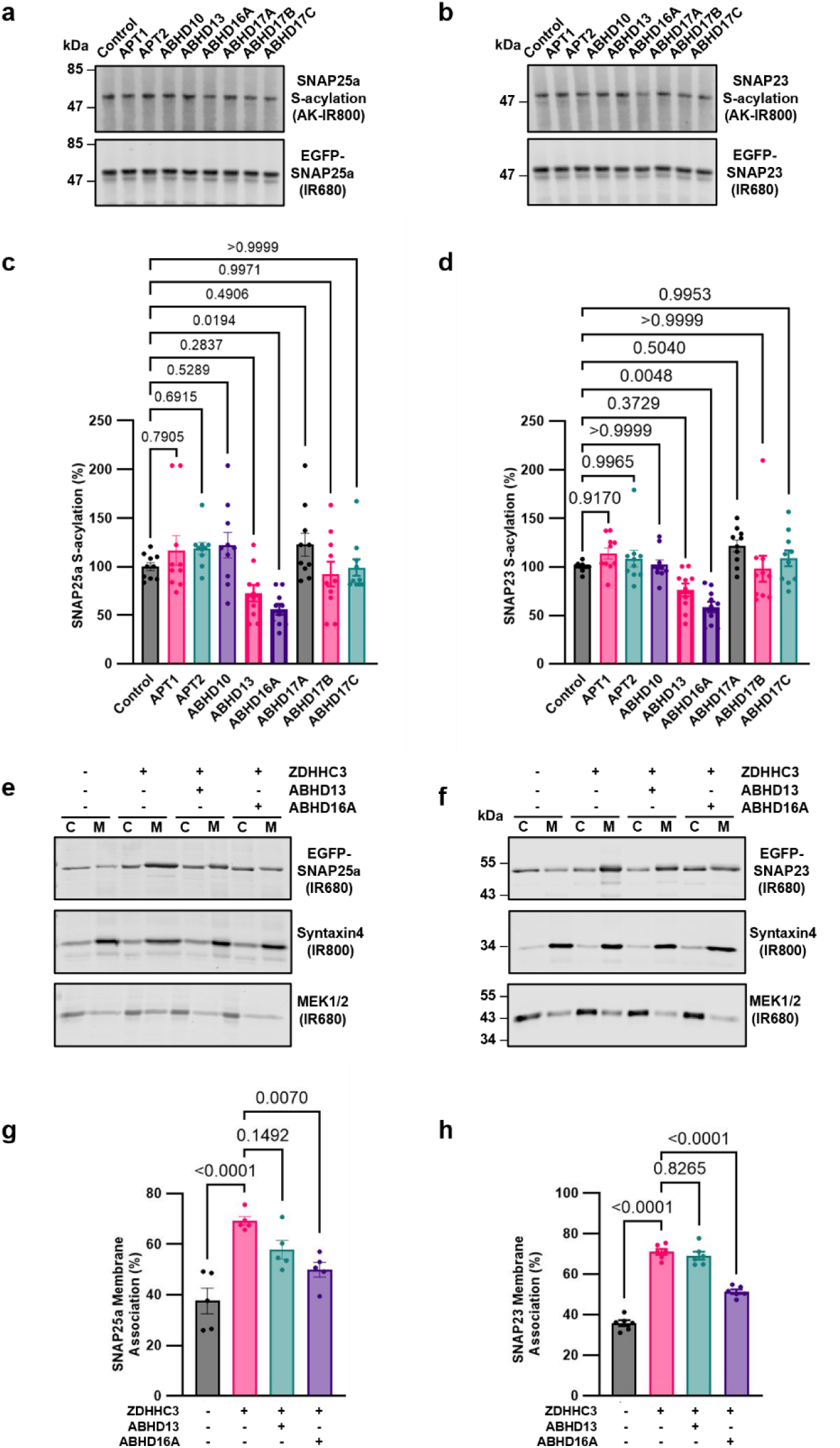
The SNAP25 isoforms SNAP25a and SNAP23 are displaced from membranes by ABHD16A-mediated deacylation only. **a., b.** *S*-acylation levels of SNAP25a (**a.**) or SNAP23 (**b.**) co-transfected with ZDHHC3 and empty vector (control) and the deacylases determined by C16:0-azide-labelling, click chemistry and immunoblotting. **c., d.** Quantification of SNAP25a **(c.)** or SNAP23 **(d.)** *S*-acylation levels shown in **a.** and **c. e., f.** Membrane partitioning of HEK293T cells transfected with SNAP25a **(e.)** or SNAP23 **(f.)** alone or with ZDHHC3 and empty vector or ABHD13 or ABHD16A. **g., h.** Quantification of SNAP25a **(g.)** or SNAP23 **(h.)** percentage membrane binding shown in **e.** and **f.** See also Supplementary Fig. 3.

### ABHD13 is selective for the first two cysteines in the SNAP25b cysteine-containing flexible linker domain

The results in Fig.3 support the idea that ABHD16A is more active towards SNAP25b than ABHD13. A possible reason for the observed difference is that ABHD16A acts on additional *S*-acylated cysteines within the SNAP25b cysteine-rich domain to those targeted by ABHD13. Previous studies have shown that reducing SNAP25b *S*-acylation sites by cysteine mutagenesis causes accumulation on recycling endosomes and the trans-Golgi network, suggesting that differential deacylation by individual thioesterases regulates SNAP25b intracellular distribution between these compartments and the plasma membrane^22^.

Therefore, we analysed whether deacylases could target specific *S*-acylated cysteines in SNAP25b. ABHD13 and ABHD16A could deacylate all SNAP25b single cysteine mutants to levels comparable to the WT protein (Supplementary Fig. 4a-d), and similar results were seen for ABHD16A towards SNAP25b double cysteine mutants (Supplementary Fig. 4e,f). However, ABHD13 deacylase activity toward the C85L/C88L mutant was significantly impaired relative to WT SNAP25b (Fig. 5a,b). These results indicate that the first two cysteines in the cysteine-rich domain of SNAP25b are required for ABHD13 deacylase activity.

**Fig. 5:**
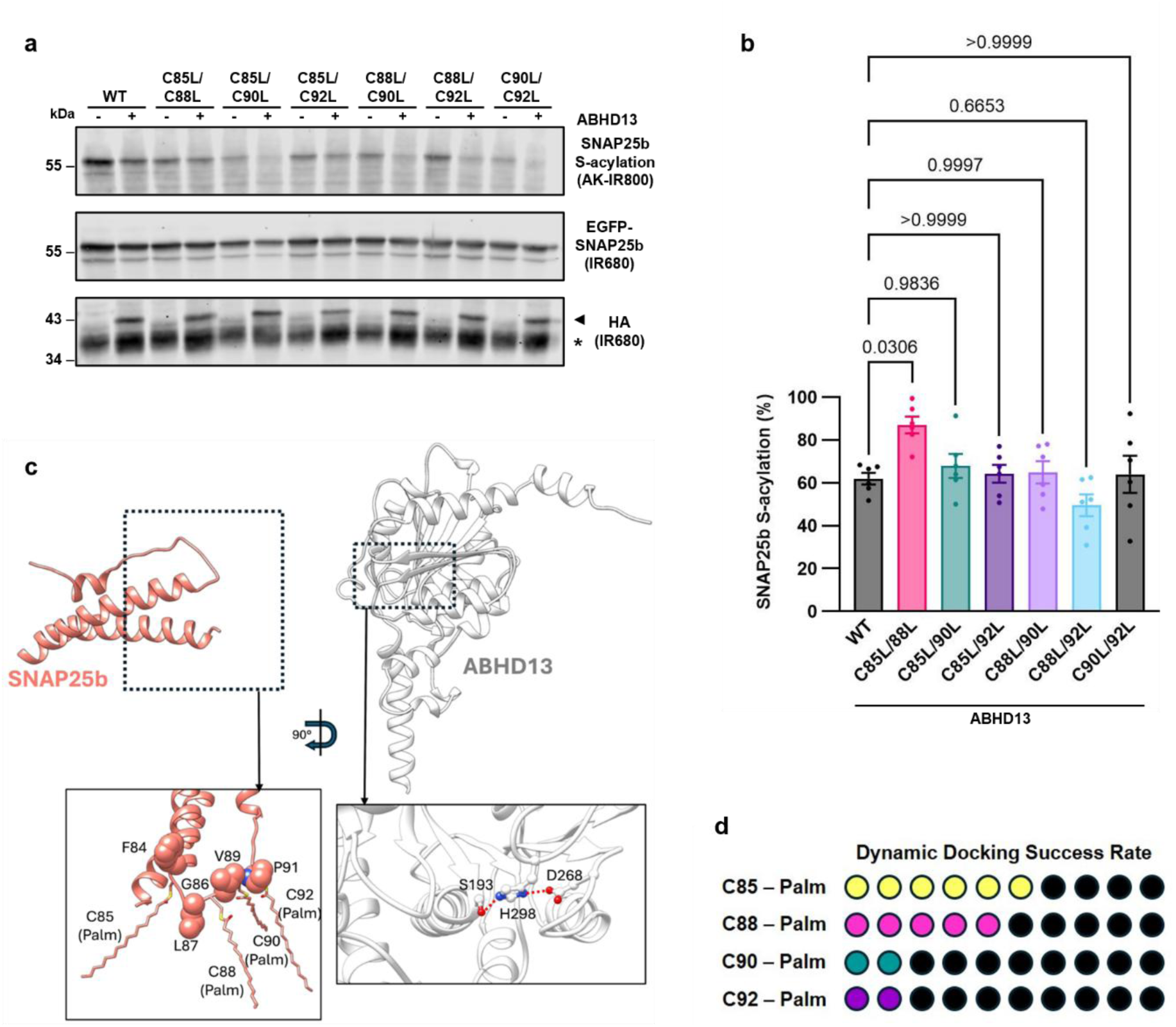
ABHD13 is selective for the first two cysteines in the cysteine-containing flexible linker domain. **a.** *S*-acylation levels of SNAP25b or C85L/C88L, C85L/C90L, C85L/C92L, C88L/C90L, C88L/C92L or C90L/C92L mutants co-transfected with ZDHHC3 and empty vector (control) or ABHD13 determined by C16:0-azide-labelling, click chemistry and immunoblotting. **b.** Quantification of SNAP25b *S*-acylation levels shown in **a. c.** SNAP25b and ABHD13 simulation setup for mwSuMD dynamic docking simulations (water molecules and ions not shown); SNAP25b residues 57-109, containing the MTD, were retained, and C85, 88, 90, and 92 were palmitoylated (left-hand insert). The ABHD13 catalytic triad is shown in the right-hand insert. **d.** Success rate of the dynamic docking mwSuMD simulations; out of 10 replicas, ABHD13 recognition of palmitoyl groups on C85 and C86 produced more stable complexes (6 and 5, respectively) than C90 and C92 palmitoyl groups (two successful replicas each).

Next, to retrieve structural information about ABHD13-SNAP25b binding and *S*-acyl group recognition, instead of performing a standard molecular docking, we used the ABHD13 AlphaFold2^37^ model retaining the cytosolic domain, and SNAP25b MTD and flanking regions (residues 57-109), to simulate with molecular dynamics (MD) their whole molecular encounter (dynamic docking) in a fully flexible and hydrated environment. This approach overcomes known limitations of molecular docking, i.e. the use of implicit solvent and conformational undersampling of highly flexible molecules^40^, like long-chain fatty acids. In addition, molecular docking, unlike MD, generally underestimates protein flexibility; this is particularly relevant here because protein side chains in AlphaFold models of deacylases may have low accuracy compared to available experimental structures (e.g. APT1^15^). Because of the dangers of using AI results unquestioningly, we successfully validated the ABHD13 AlphaFold2 model using a multiple template sequence alignment method known to work within the twilight zone^41^, with ABHD13 structural homologues retrieved from the Dali protein comparison webserver^42^ – see the associated article^43^.

Starting from the fully dissociated ABHD13 – SNAP25b complex (Fig. 5c), the adaptive sampling method supervised MD (mwSuMD^44^) was used for simulating 40 binding events (i.e. 40 dynamic dockings carried out on 40 separate mwSuMD replicas) of the simplified system formed by the ABHD13 αβ cytosolic domain and SNAP25b. Ten dynamic dockings for each *S*-palmitoylated cysteine (i.e. C85, C88, C90, and C92) were sampled by supervising the distance between the palmitoyl carbonyl group and S193 (Video S1). As palmitoylation is the most prevalent type of *S*-acylation, our simulations were built with palmitic acid as the modifying fatty acid. We considered a dynamic docking productive if the distance between the carbonyl carbon atom of the palmitoylated cysteine and the oxygen atom of the catalytic S193 reached and remained below 10 Å for the whole duration of the simulation. The dynamic docking success rate of C85 and C88 (six and five, respectively), expressed as the number of productive simulations out of 10, was higher than C90 and C92 (two successful simulations each) (Fig. 5d). This suggests that the *S*-palmitoyl group of C85 and C88 have better complementarity for ABHD13, and qualitatively supports the experimental evidence that the first two cysteines in the cysteine-rich domain of SNAP25b are more important for ABHD13 deacylase activity than the other two (Fig. 5a,b).

### Revealing the selectivity of ABHD13 towards SNAP25 isoforms

To determine why ABHD13 prefers SNAP25b over SNAP25a and SNAP23, we aligned their primary sequences and found one single amino acid shared between SNAP25a and SNAP23, C84 and C79, respectively, whereas SNAP25b has a phenylalanine (F84) at the same position (Supplementary Fig. 1 and Fig. 1b). Since ABHD13 preferentially reacts with *S*-acylated cysteines at position 85 and 88 of SNAP25b (Fig. 5), we hypothesised that it is not the presence of a phenylalanine at position 84 that promotes ABHD13-mediated deacylation of SNAP25b, but rather an additional *S*-acylated cysteine immediately upstream of C85 may limit the ability of ABHD13 to access these acyl chains (Fig. 6a). We tested the ability of ABHD13 or ABHD16A to deacylate SNAP25b F84A and F84C mutants (Fig. 6b,c) and SNAP25a and SNAP23 mutants with reversed cysteine/phenylalanine at this position (C84F and C79F, respectively; Fig. 6d-g). Replacing phenylalanine at position 84 in SNAP25b with cysteine, therefore mimicking the configuration of SNAP25a and SNAP23, resulted in a loss of deacylation activity by ABHD13 compared to WT SNAP25b (Fig. 6b,c). Conversely, mutation of the first cysteine in SNAP25a (C84F) and SNAP23 (C79F), mimicking the phenylalanine-cysteine configuration of the SNAP25b cysteine-rich domain, led to a significant increase in deacylation by ABHD13 of these isoforms (Fig. 6d-g). Importantly, introducing an alanine residue in SNAP25b at position F84 (F84A) did not decrease ABHD13-mediated deacylase activity (Fig. 6b,c). Although we observed a slight increase in ABHD16A-mediated deacylation of the SNAP25b (F84C) mutant, there was no difference between SNAP25a (C84F) and SNAP23 (C79F) compared to the wild-type protein (Fig. 6d-g). We repeated the dynamic docking shown in Fig. 5, this time modelling SNAP25b (F84C) *S*-palmitoylated C85 in the co-presence of *S*-palmitoylated C84. In these simulated conditions, the success rate of C85 depalmitoylation reduced from six to two dynamic docking events, indicating that physical separation between C84 and C85 was required for productive engagement of ABHD13 (Supplementary Fig. 5). Such molecular behaviour likely explains the ABHD13 selectivity towards SNAP25b and further supports the hypothesis that tandem cysteines at the N terminus of the cysteine-rich region hinders ABHD13 activity. Collectively, these results show that the position and arrangement of *S*-acylated cysteines are an important determinant for ABHD13 deacylase activity.

**Fig. 6:**
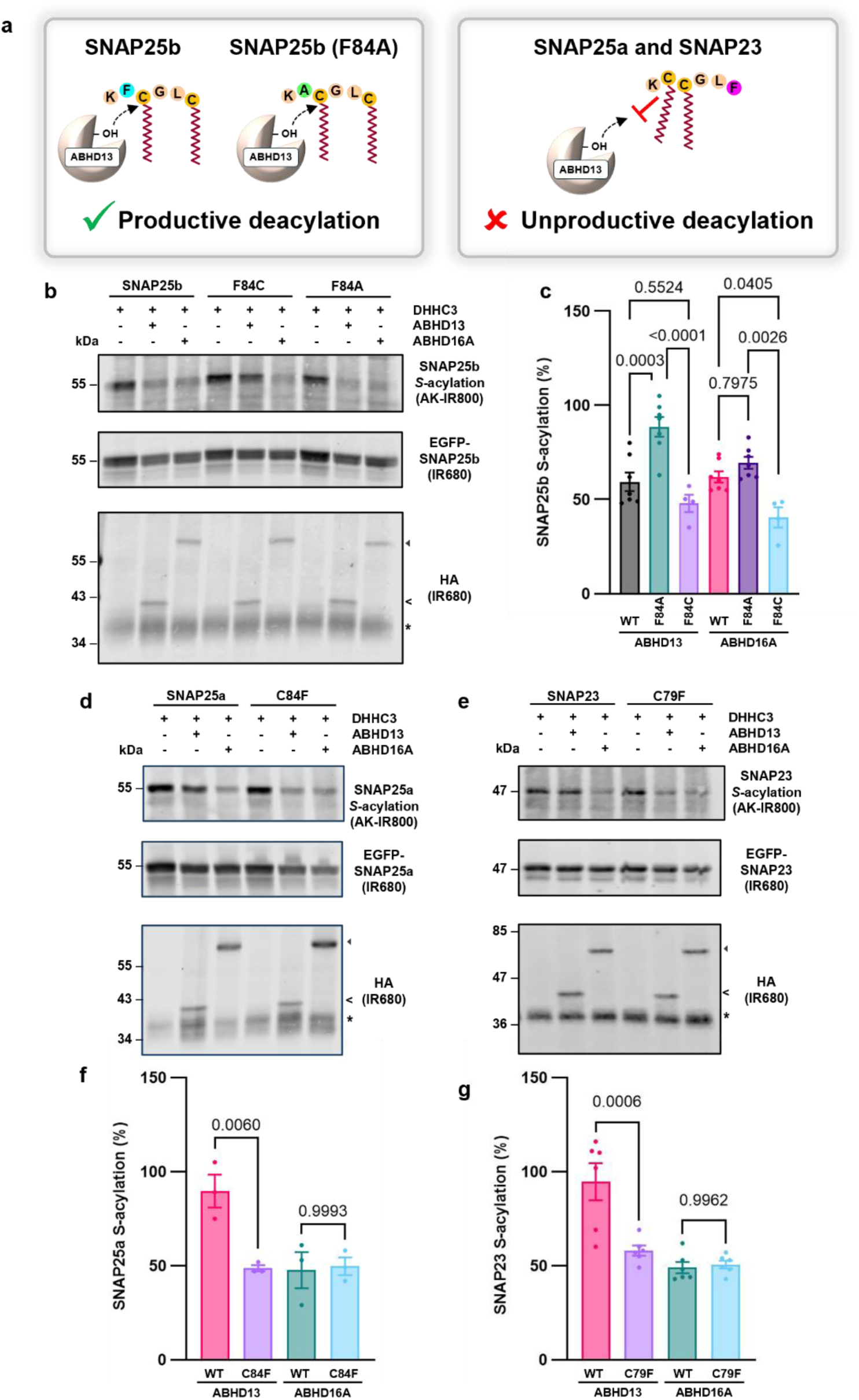
Two consecutive *S*-acylatable cysteines inhibit ABHD13-mediated deacylation. **a.** Schematic of productive *versus* unproductive ABHD13-mediated deacylation towards SNAP25 mutants and isoforms. **b., d., e.** *S*-acylation levels of SNAP25b WT, F84C, or F84A **(b.)**, SNAP25a WT or C84F **(d.)**, or SNAP23 WT or C79F **(e.)** mutants co-transfected with ZDHHC3 and empty vector (control) or ABHD13 determined by C16:0-azide-labelling, click chemistry and immunoblotting. **c., f., g.** Quantification of *S*-acylation levels shown in **b, d., e.**

### Identification of a hydrophobic pocket and key residues regulating ABHD13 deacylase activity

To investigate further the mechanisms that could dictate ABHD13 activity, we sought to understand the structural characteristics of ABHD13. Lipid binding pockets have been identified for many mSH deacylases, including ABHD17 and APT2^11,12,15^. However, whether ABHD13 contains such a pocket is unknown, as are the mechanisms that regulate its deacylase activity. We analysed the mwSuMD dynamic docking simulations to determine the preferred positions occupied by the aliphatic chain of *S*-palmitoyl groups. Interestingly, all four palmitoyl groups explored a hydrophobic pocket located below the catalytic site (Fig. 7a) whose accessibility appeared regulated by residues L65, I222 (both located in the proximity of S193), and L229 (placed slightly more distant to the catalytic triad). To examine the role of these hydrophobic residues for acyl chain binding, we created individual tryptophan or aspartate mutants at positions L65, I222 and L229 and assessed the effect on the incorporation of C16:0-azide on SNAP25b. Mutation of L65 to either D or W, or I222 to W (I222D did not express), resulted in a complete loss of ABHD13 deacylase activity towards SNAP25b (Fig. 7b,c), suggesting that these amino acids are essential for enzymatic activity. Interestingly, whilst the L229D mutant prevented ABHD13-mediated SNAP25b deacylation, the L229W mutant did not, suggesting that a leucine at position 229 is not required for deacylase activity, *per se*, but rather the integrity of the putative lipid binding pocket depends on a hydrophobic residue at this position (Fig. 7b,c).

**Fig. 7:**
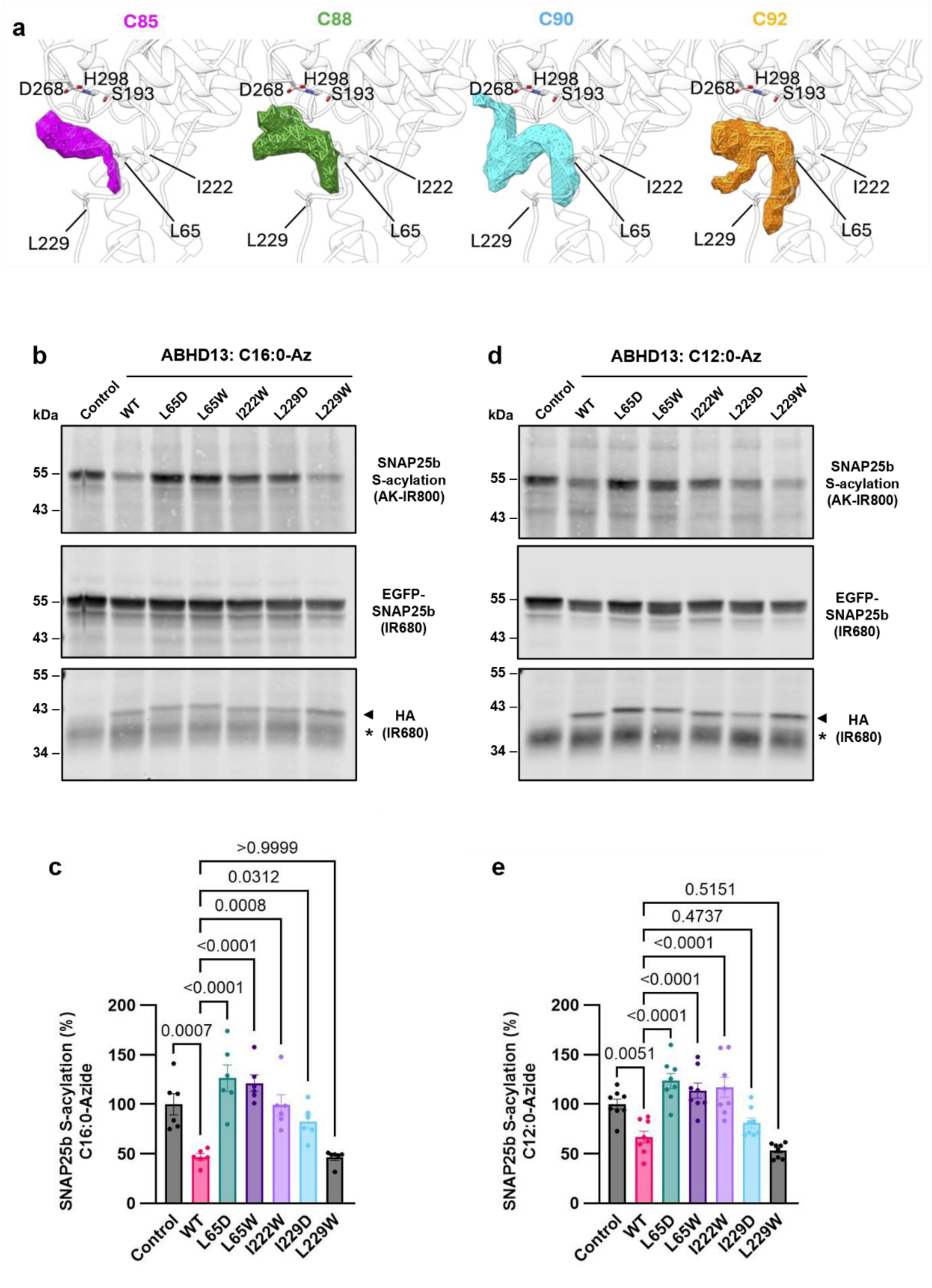
Identification of a hydrophobic pocket in ABHD13. **a.** Volumetric maps (probability > 20%) of the palmitoyl groups during mwSuMD of the simplified system. ABHD13 is shown as transparent white ribbon; the position of the catalytic residues S193, S268, and H298, and of the important hydrophobic residues is indicated. Volumes occupied by C85, C88, C90, and C92 palmitoyl group atoms are coloured in magenta, green, cyan, and orange, respectively. **b., d.** *S*-acylation levels of SNAP25b co-transfected with ZDHHC3 and empty vector (control), ABHD13 WT or pocket mutants determined by C16:0 **(b)** or C12:0 **(d)** - azide-labelling, click chemistry and immunoblotting. **c.** Quantification of SNAP25b *S*-acylation levels shown in **b.** and **d.**

### A complete model of ABHD13- SNAP25b recognition

We exploited the structural insights reported above to build a complete model of the complex between ABHD13 and SNAP25b (*S*-palmitoylated at C85, C88, C90, and C92), positioning both structures relative to the membrane using the Positioning of Protein in Membrane server 2.0^45^ (Fig. 8, Supplementary Fig. 6, Video S2). After preparation for MD simulation, we performed three replicas of mwSuMD^44^, sampling the approach of the *S*-palmitoylated SNAP25b C85 carbonyl carbon atom to the ABHD13 S193 oxygen atom to study the structural events preceding and favouring deacylation. Out of three mwSuMD replicas (Fig. 8b), two reached less than 5 Å (minimum distance = 3.7 Å), indicating that proximity between substrate and catalytic serine is consistent, or very close, to that required for the reaction to take place based on hydrolysis predictions (i.e. ∼3.5 Å)^46^. Overall, SNAP25b swings with an angle comprised between ∼45° and 60° to the membrane plane, while the ABHD13 membrane-interacting helix at ∼60°, with the amphipathic N-terminal helix positioned at the interface between the membrane and the inner side of the simulation box (Fig. 8a). Interestingly, ABHD13 residues 220-245 formed a U-shaped element that spanned the membrane and deformed the inner side, as shown by the position of the phospholipid heteroatoms in Fig. 8a, and increased cholesterol local concentrations through direct interactions (Fig. 8a, insertion). Early during the equilibration stage, the *S*-palmitoyl group of C85 spontaneously inserted its terminal half into the hydrophobic pocket shaped by L65, I222, and L229 directly from the membrane. During the following mwSuMD simulations, the thioester group was presented to the catalytic triad just above the phospholipid polar heads, but keeping the aliphatic chain buried within the lipophilic environment formed by the ABHD13 hydrophobic tunnel (Fig. 8b, Video S2). The rapid insertion of the terminal part of the *S*-palmitoyl group of C85 into the ABHD13 hydrophobic pocket raises the question of whether aliphatic chains shorter than palmitate can be accepted by ABHD13 with different efficiency.

**Fig. 8:**
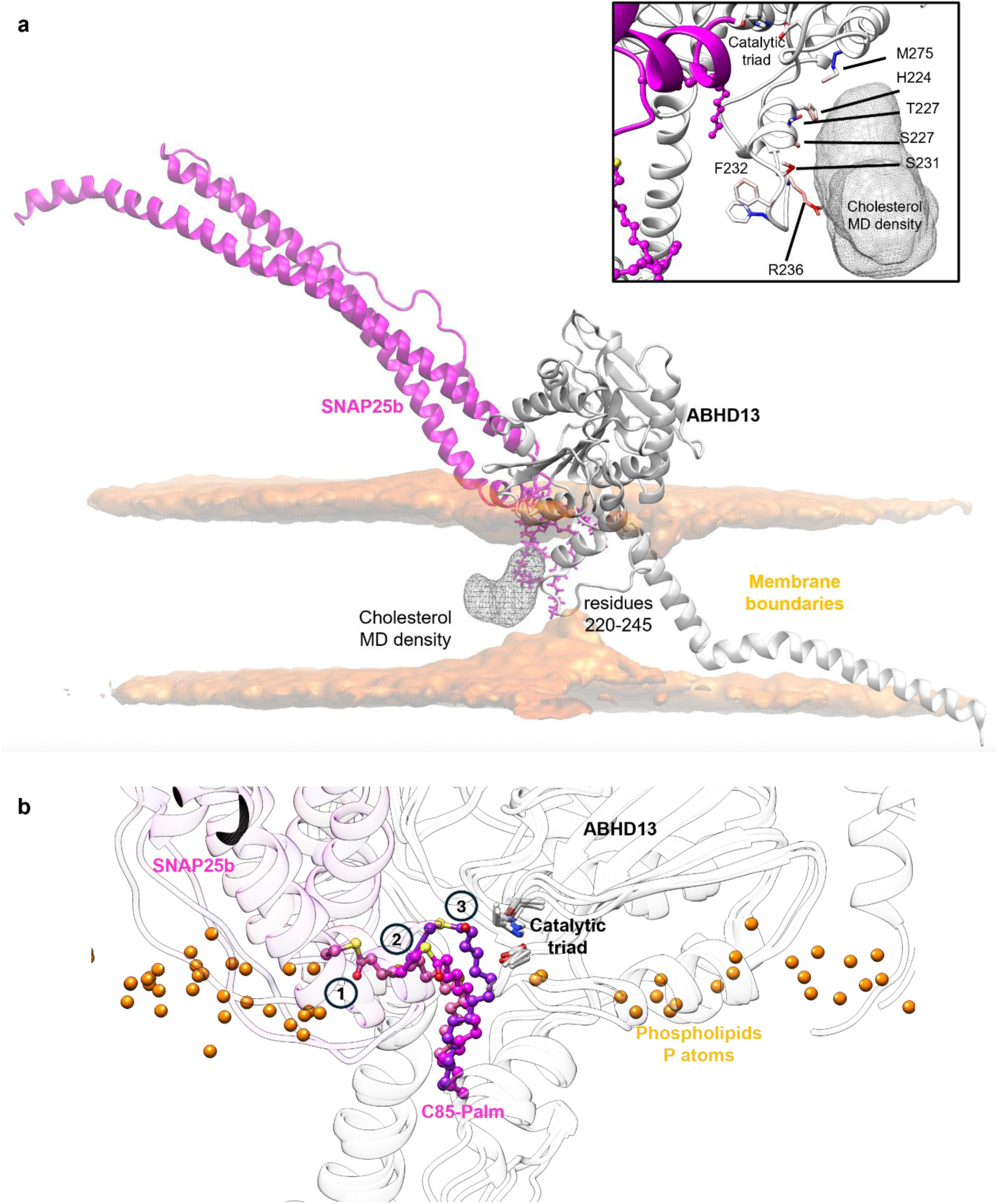
SNAP25 – ABHD13 interactions during mwSuMD. **a.** Volumetric maps of the membrane polar atoms (orange, occupancy > 20%) and cholesterol molecules (dashed grey, occupancy > 10%) during mwSuMD simulations, overlaid on the SNAP25:ABHD13 initial complex. Insertion: details of ABHD13 residues that interacted with cholesterol during mwSuMD simulations; side chain atoms are coloured from blue (no interactions) to red (maximum interaction). **b.** SNAP25b C85 palmitoyl group (pink to purple) inserted into ABHD13 hydrophobic tunnel directly from the membrane; three mwSuMD snapshots show the path of the C85 palmitoyl group, with phospholipid P-atoms represented as orange spheres. The catalytic triad side chains (S193, H298, and E268) are shown as white sticks for reference.

We experimentally tested ABHD13 thioesterase activity on SNAP25b *S*-acylated with azido-fatty acids containing shorter (lauric, C12:0 and myristic, C14:0) or longer (stearic; C18:0) aliphatic chains compared to palmitic acid, or with a perturbed (unsaturated) stereochemistry (Δ9-oleic acid-azide, C18:1). All azido-fatty acids tested were fully functional substrates of ABHD13 (Fig. 7b-e and Supplementary Fig. 7). Since the proximity of the ABHD13 L65, I222 and L229 mutants to the catalytic triad of ABHD13 seems related to its thioesterase activity, we further examined the ability of these mutants to accept C12:0-azide and C14:0-azide. Whilst the ABHD13 mutants had reduced activity towards C14:0-azide, consistent with those observed for the longer *S*-palmitoyl group, the shorter C12:0-azide was an effective substrate for both ABHD13 L229 mutants (Fig. 7d,e and Supplementary Fig. 7). This chain length-dependent activity of ABHD13 mutants suggests that an intact hydrophobic pocket is necessary only for longer chain substrates. Indeed, the palmitoyl terminal carbon atoms C14-C16 were the most engaged in hydrophobic interactions during simulations (Supplementary Fig. 8), indicating higher dependence on the hydrophobic pocket for the recognition of long-chain fatty acids than for short ones, which could be extracted from the membrane and stabilised by fewer contacts. This scenario is consistent with a slightly divergent recognition mechanism driven by the different substrate chain lengths, and it is supported by desorption experiments of fatty acids from phospholipid vesicles, where shorter fatty acids required less energy to be extracted^47^.

## DISCUSSION

Altered *S*-acylation dynamics or dysregulated deacylase activity are implicated across many neurological, neurodegenerative, developmental and psychiatric disorders, but emerging evidence indicates that manipulating deacylase activity is a promising therapeutic strategy in Alzheimer’s, Parkinson’s, and Huntington’s disease models^48^. Outside the nervous system, inhibiting the deacylation machinery is an effective approach for treating acute myeloid leukaemia and inflammatory bowel disease^49–52^. Despite the abundance of *S*-acylated proteins and the physiological importance of the modification, no consensus sequences define *S*-acylation or deacylation^1^. Deacylases discriminate among highly similar lipid-modified cysteines across diverse substrates. For example, APT1 specifically deacylates H-Ras, but not other Ras isoforms, while ABHD17 regulates N-Ras and PSD-95^16,28,53^ ^13,53,54^. APT1 and LYPLAL1 – but not APT2 – target two distinct *S*-acylated domains within BK channels, yet ABHD17A selectively deacylates only the STREX domain, enabling splice variant–specific regulation^29,55^.

Here, we provide evidence of a new example of biased deacylation, with ABHD13 and ABHD16A showing distinct selectivity towards SNAP25 isoforms. The isoforms a and b differ by 9 amino acids and are differentially expressed: SNAP25a during embryonic and early development, and SNAP25b predominantly in postnatal and adult brain^30^. SNAP25a and SNAP23, a ubiquitously expressed homologue, have a second *S*-acylated cysteine immediately upstream of C85. We found that replacing this cysteine in SNAP25a (C84) or SNAP23 (C79) with alanine or phenylalanine, the latter of which mimics the *S*-acylation domain present in SNAP25b, restores ABHD13-mediated deacylation of these proteins, whereas a SNAP25b mutant containing the additional cysteine (F84C) disrupts ABHD13 activity (Fig. 6). Our adaptive sampling MD simulations identified a hydrophobic pocket in ABHD13 that is essential for activity (Fig. 7), and suggested that two adjacent *S*-acylation sites compete for binding, reducing productive substrate engagement and explaining the selectivity of ABHD13 towards distinct SNAP25 isoforms (Supplementary Fig.5). These findings identify molecular determinants of ABHD13 activity toward SNAP25b, and support two previous studies: one showing a double cysteine motif in GAP-43 favours APT2 over APT1 ^56^, and another showing the deacylation rate of recombinant APT1/2 towards synthetic *S*-acylated peptides is influenced by the chemical nature of the amino acids surrounding the modified cysteine *in vitro* ^27^. Collectively, these observations strongly suggest that cysteine configuration influences deacylase activity and substrate selectivity. Furthermore, we found that ABHD13 preferentially deacylates the two N-terminal *S*-acylation sites on SNAP25b (C85 and C88; Fig. 5). Interestingly, a previous study showed that mutating these two cysteines to leucine (C85L/C88L) increased SNAP25b localisation to recycling endosomes and the trans-Golgi network, implying that ABHD13-mediated site-specific deacylation may regulate SNAP25b intracellular targeting^22^. The number of *S*-acylated cysteines has also been linked to the presence of SNAP25 in cholesterol microdomains and its exocytotic capacity at the plasma membrane ^20,21^.

We identified a hydrophobic pocket in ABHD13 and found that the orientation of the side chains of the hydrophobic residues form a cavity that regulate ABHD13 activity (Fig.7). Mutation of residues L65, I222 and L229 (unless mutated to tryptophan) that form the hydrophobic pocket (Fig. 7b,c, 8b, Video S2) led to a loss of ABHD13 activity, indicating the structural integrity of the region that accommodates the fatty acids is a determinant factor for enzymatic activity. These results are in agreement with previous observations on the structure of APT1 and APT2 that revealed a hydrophobic pocket is essential for their activity ^15^, with mutations within the region in APT2 leading to a reduction in enzymatic activity ^11^.

Based on MD simulations performed by Abrami *et al.,* membrane deformation was proposed as the mechanism of action of the peripheral deacylase APT2 due to an increase in membrane thickness around the acyl group^11^. Such deformation is supposed to favour substrate extraction from the membrane towards a shallow hydrophobic binding site located horizontally just above the membrane (Fig. 9a)^11^. In our mwSuMD simulations, ABHD13 recognised the SNAP25b palmitoylate *via* a different mechanism (Fig. 9b), in which the hydrophobic tail is only partially extracted from the lipophilic membrane environment, owing to the protection provided by the hydrophobic pocket beneath the catalytic site (lined by L65, I222, and L229). We did not observe a membrane thickness increase analogous to APT2, but rather a local decrease in thickness of the leaflet opposite to the acyl chain cleaved, due to the partially hydrophilic nature of a U-shaped tongue that spans the membrane (ABHD13 residues 220-245). The local membrane deformation was accompanied by an increase in cholesterol occupancy nearby ABHD13. Although speculative, it is possible that membrane deformation and local rigidification, due to higher cholesterol concentration, favour the *S*-acyl group recognition by tuning the dynamics of phospholipids near ABHD13. Since SNAP25b is a substrate for ABHD13, the higher cholesterol concentration seen around ABHD13 is consistent with the association between the extent of SNAP25 *S*-acylation and its partitioning into cholesterol-rich membrane microdomains in neuroendocrine cells^20,21^.

**Fig. 9:**
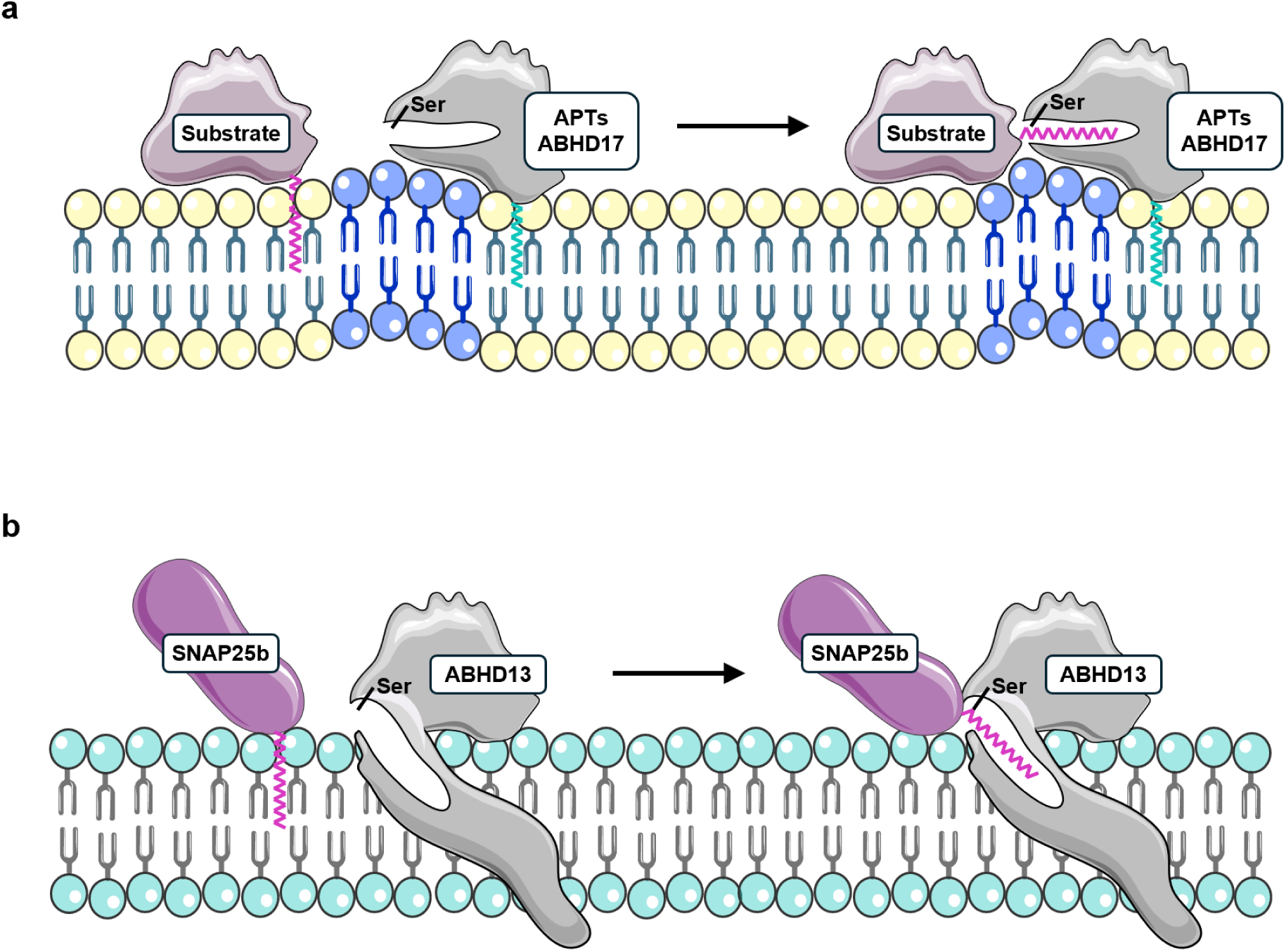
Proposed model for SNAP25b acyl group recognition by ABHD13. **a.** In APT1, APT2, and ABHD17, membrane deformation favours the extraction of the substrate acyl group from the membrane towards a shallow horizontal hydrophobic pocket located just above the membrane. **b.** In ABHD13, the SNAP25b acyl group is only partially extracted from the membrane into the hydrophobic pocket located underneath the catalytic site.

Whilst APT1 and APT2 depend on reversible *S*-acylation for stable membrane attachment and hence their activity^53^, ABHD13 and ABHD16A contain hydrophobic helices that can facilitate membrane binding. Interestingly, ABHD17 is thought to be peripherally membrane-associated through non-reversible *S*-acylation^12^, suggesting that these ABHD isoforms could represent a new class of membrane deacylases with alternative mechanisms for regulating their activity independently of reversible membrane binding. Whilst the subcellular localisation of ABHD13 has not been determined, ABHD16A is found on the cytoplasmic face of the cellular membranes^4^, including the plasma membrane of human platelets and mouse megakaryocytes^57^. However, it is also found on the ER in mouse brain and Neuro-2a cells^58^, and at ER-mitochondrial contact sites in HeLa cells, osteosarcoma (U2OS) cells and mouse embryonic fibroblasts (MEFs), where it regulates mitochondrial fission/fusion by modulating local phospholipid composition^59^. How might an ER-resident ABHD16A deacylate SNAP25, which is initially *S*-acylated by Golgi-localised ZDHHC proteins for plasma membrane targeting^23^? We propose that such interactions can occur at membrane contact sites (MCS), which act as central hubs for intracellular communication and protein regulation across multiple compartments^60^. This would enable membrane-associated deacylases to rapidly and locally regulate *S*-acylation dynamics, similar to how the peripheral membrane deacylases APT1 and APT2 control protein localisation and function by shuttling between the cytosol and membranes through reversible *S*-acylation.

It has been suggested that the overexpression of deacylases may drive non-physiological or stochastic substrate engagement, so caution is needed when interpreting overexpression assays to ensure activity reflects intrinsic substrate selectivity^10^. In addition to our findings on ABHD13 substrate selectivity, we found that while ABHD16A efficiently deacylates all SNAP25 isoforms (Fig. 4a–d), it fails to deacylate dually *S*-acylated H-Ras (Supplementary Fig. 2f,g). This indicates that ABHD16A can discriminate between substrates, even though overall it appears more active than ABHD13 under the same conditions (Fig. 3a–d), indicating that catalytic action is not indiscriminately enhanced simply by increased expression. This conclusion aligns with Yokoi *et al.* (2019), who overexpressed ABHD16A yet saw no deacylation of PSD-95^16^. Thus, although overexpression remains an important technical caveat, structural and compartmental constraints may preserve substrate specificity even under elevated enzyme levels.

Our pulse-chase analysis determined that THL could preserve SNAP25 deacylation to a similar extent as PalmB in HEK293T cells (Fig. 2a,b), but we cannot rule out other endogenously expressed thioesterases influencing SNAP25 *S*-acylation dynamics. The development of selective cell-permeable ABHD13 and ABHD16A inhibitors, supported by structural determination, along with selective knockdown or knockout of deacylases in cells and tissues expressing SNAP25, will be necessary to better understand the physiological role of these enzymes. Furthermore, future efforts should focus on characterising the novel deacylases, including ABHD13, ABHD16A, and ABHD17, using purified recombinant proteins to measure reaction kinetics, since only APT1/2 and PPT1 deacylase activity has been confirmed *in vitro*.

Our findings explain how ABHD13 distinguishes between two highly similar *S*-acylated substrates - SNAP25a and SNAP25b - and between distinct *S*-acylation sites within the same protein, which may underlie the differential regulation of the two isoforms *in vivo*. *S*-acylation is essential for neuronal development, neurotransmission and synaptic plasticity, and changes in SNAP25 are linked to metabolic diseases and a wide spectrum of neurological and psychiatric diseases, including ADHD, schizophrenia, bipolar disorder, epilepsy, intellectual disability, Alzheimer’s and Parkinson’s disease. Our work identifying the SNAP25 deacylases not only clarifies fundamental synaptic biology but opens new avenues for understanding and potentially targeting mechanisms that may underlie synaptic dysfunction in neurodevelopmental and neurodegenerative disorders.

## ACKNOWLEDGEMENTS

We thank Dr Christine Salaun (University of Strathclyde, UK) and Dr Liam Butler (Glasgow Caledonian University, UK) for advice on the GFP-Trap. We thank Prof Masaki Fukata (National Institute for Physiological Sciences, Japan) for providing the pEF-BOS-HA-tagged ZDHHC plasmids. This work was supported by UKRI [Future Leaders Fellowship MR/W011840/1], the Academy of Medical Sciences [Springboard award SBF005\1122] and Tenovus Scotland [Grant S16/11] to J.G. We are grateful to the Centre for Health and Life Sciences and the Doctoral College at Coventry University for stipend and fee support for studentships to L.M. and C.N.

## AUTHOR CONTRIBUTIONS

Conceptualisation, L.M., L.P., C.N., N.C.O.T., C.A.R., G.D. and J.G.; formal analysis, L.M., L.P., C.N., C.A.R., G.D. and J.G.; funding acquisition, J.G.; investigation, L.M., L.P., C.N., C.A.R., G.D. and J.G.; methodology, L.M., L.P., C.N., C.A.R., G.D. and J.G.; project administration, J.G.; resources, N.C.O.T.; software; supervision, J.G., C.A.R., and G.D.; writing–original draft, L.M., L.P., G.D., and J.G.; writing–review & editing, L.M., L.P., C.N., N.C.O.T., C.A.R., G.D., and J.G.

## MATERIALS AND METHODS

### Plasmids and molecular biology

pEF-BOS-HA and the HA-ZDHHC plasmids were kindly provided by Professor Masaki Fukata (National Institute for Physiological Sciences, Japan). pEGFP-C2, EGFP-SNAP25b, EGFP-SNAP25a, EGFP-SNAP23 and their mutants were as previously described^23^. EGFP-HRas subcloned into pEGFP-C3^61^ and murine ABHD16A subcloned into pEF-BOS-HA were described previously^62^. cDNA encoding human APT1 (NM_006330.4), APT2 (NM_007260.3), ABHD10 (NM_018394.4), ABHD13 (NM_032859.3), ABHD17A (NM_031213.4), ABHD17B (NM_016014.4), ABHD17C (NM_021214.2), and mouse ABHD6 (NM_025341.4) were synthesised by Thermo Fisher Scientific and sub-cloned into pEF-BOS HA. ABHD16A(D430A) subcloned into pEF-BOS-HA and EGFP-tagged SNAP25b (F84W, F84Y, F84A) SNAP25a (C84W, C84A) and SNAP23 (C79W, C79Y, C79A) were generated by PCR using a QuikChange II XL Site-Directed Mutagenesis Kit. ABHD13(D268A) subcloned into pEF-BOS-HA was synthesised by Genscript. The validity of all constructed plasmids was confirmed by Sanger sequencing of both strands by Eurofins Genomics.

### Antibodies

The Rat monoclonal anti-HA (RRID:AB_390918) was from Roche. A Living Colors A.v. Monoclonal Antibody against GFP (JL-8; RRID:AB_10013427) was from Takara Bio. The recombinant Anti-Syntaxin 4 antibody (RRID:AB_2891056) was from Abcam. A MEK1/2 (D1A5) Rabbit monoclonal antibody (RRID:AB_10829473) was from Cell Signaling Technology. The IRDye 680RD Goat anti-Rat (RRID:AB_2814913), IRDye 680RD Donkey anti-Mouse (RRID:AB_2814912), IRDye 680RD Donkey anti-Rabbit (RRID:AB_10954442) and IRDye 800CW Donkey anti-Rabbit (RRID:AB_621848) were from LI-COR Biosciences.

### Cell culture and transfections

Human Embryonic Kidney 293T cells (HEK293T; RRID:CVCL_0063) purchased from the European Collection of Authenticated Cell Cultures (ECACC) were cultured in high glucose Dulbecco’s Modified Eagle Medium (DMEM-GlutaMAX®, Gibco) supplemented with 10% heat inactivated foetal bovine serum (FBS, Gibco) at 37°C in a humidified atmosphere containing 5% CO_2_. Cells were routinely tested as mycoplasma negative using a Lonza™ MycoAlert™ Mycoplasma Detection Kit (Fisher Scientific). Cells were transiently transfected using Lipofectamine 2000 reagent (Invitrogen) according to manufacturer instructions and harvested the following day.

### Bioorthogonal labelling, Click chemistry and pulse-chase analysis

Transfected HEK293T cells growing in 24-well plates were washed once with 500 μl of warm PBS and incubated in 300 μl/well of serum-free DMEM supplemented with 1 mg/mL defatted BSA (Merck Life Science Ltd, UK) and 100 µM of fatty acid acid-azide (C12:0-azide, C14:0-azide, C16:0 azide and C18:0-azide were synthesised as before^26^; C18:1-azide was purchased from Avanti Research) for 4 hours at 37°C in a humidified atmosphere containing 5% CO_2_. Fatty acid acid-azide-labelled cells were washed twice in cold PBS then lysed on ice in 100 μL of lysis buffer (50 mM Tris pH 8.0, 0.5% (w/v) SDS) containing protease inhibitors. Conjugation of alkyne IR-800 Dye (AK-IR800) (LI-COR) to fatty acid acid-azide was carried out for 1 h at room temperature with end-over-end rotation by adding 80 μL of fresh click chemistry reaction mixture (2.5 μM AK-IR800, 4 mM CuSO_4_, 400 μM TBTA (Tris[(1-benzyl-1H-1,2,3-triazole-4yl) methyl]), and 20 μL of 40 mM ascorbic acid. The reaction was terminated by the addition of 67 µL of 4 x SDS loading buffer (6 mM bromophenol blue, 200 mM Tris pH 6.8, 40% (w/v) glycerol, 8% (w/v) SDS) containing 100 mM dithiothreitol (DTT). The samples were incubated at 95 °C for 5 min, separated by SDS-PAGE, transferred to nitrocellulose for immunoblotting analysis, and visualised using a LI-COR Odyssey Imaging System.

For pulse-chase analysis, transfected HEK293T cells growing in duplicate 24-well plates were labelled with C16:0-azide as before. After 4 hours, one plate (the ‘pulse’) was processed for the click chemistry reaction, whilst the other plate (the ‘chase’) was washed once in warm DMEM and incubated for a further 3 hours at 37°C in a humidified atmosphere containing 5% CO_2_ in 350 μl/well of serum-free DMEM supplemented with 1 mg/mL fatty acid free BSA, 500 µM of palmitic acid (Merck), 50 µg/mL cycloheximide and 1%(v/v) dimethyl sulfoxide (DMSO; Control), 100 µM Palmostatin B (Calbiochem), 10 µM WWL70 (Cayman Chemical), or 100 nM tetrahydrolipstatin (THL; Merck) prior to click chemistry and immunoblot analysis, as before.

### Co-immunoprecipitation

Transfected HEK293T cells growing in 6-well plates were scraped into 200 µL of lysis buffer (PBS containing 0.5 % Triton X-100 and protease inhibitors) and incubated for 30 minutes on ice. Cell lysates were centrifuged at 14,000 x *g* for 5 minutes at 4 °C and the supernatant was adjusted to 500 µL using ice-cold PBS. Fifty microliters were saved as the ‘input’ and the remainder was mixed with 10 µL bed volume of activated ChromoTek GFP-Trap beads (Proteintech) and incubated for 1 hour with end-over-end rotation at 4 °C. Beads were recovered by centrifugation and washed in PBS. Fifty microliters of 2 × SDS loading buffer containing 50 mM DTT was added, and bound proteins were eluted by incubating the beads at 95 °C for 10 minutes. Supernatants containing the Bound samples were collected and subjected to SDS-PAGE, transferred to nitrocellulose for immunoblotting analysis, and visualised using a LI-COR Odyssey Imaging System.

### Cell fractionation

The fractionation protocol was adapted from Baghirova *et al* ^63^. Transfected HEK293T cells growing in 12-well plates were scraped into 500 µL of cold PBS; 10 % was saved as the ‘input’. The remaining cells were centrifuged at 500 x *g* for 10 minutes at 4 °C and resuspended in 150 µL of cold lysis buffer A (150 mM NaCl, 50 mM HEPES pH 7.4, 25 µg/mL digitonin, 1 M hexylene glycol) containing protease inhibitors and incubated on ice for 10 minutes, followed by centrifugation at 2000 x *g* for 10 minutes at 4 °C. The supernatant containing the cytosolic fraction was reserved and the pellet was incubated in 150 µL of cold lysis buffer B (150 mM NaCl, 50 mM HEPES pH 7.4, 1% (v/v) Igepal, 1 M hexylene glycol) containing protease inhibitors for 30 minutes on ice. Insoluble material was eliminated by centrifugation at 7000 x *g* for 10 minutes at 4 °C and the supernatant containing the membrane fraction was collected. All fractions (input, cytosolic and membrane) were supplemented with 4 x SDS loading buffer containing 100 mM DTT and incubated at 95 °C for 5 min. Equal volumes of the cytosolic and membrane fractions were subjected to SDS-PAGE, transferred to nitrocellulose for immunoblotting analysis, and visualised using a LI-COR Odyssey Imaging System.

### Membrane isolation and Fluorophosphonate-azide activity profiling

Transfected HEK293T cells in 10cm^2^ dishes were scraped into ice-cold PBS, passed 20 X through a 25G needle and centrifuged at 8000 x *g* for 10 minutes at 4 °C. The supernatant was then passed 20 X through a 25G needle and centrifuged at 100,000 × *g* for 1 hour at 4 °C and the pellet containing isolated membranes was resuspended in 500 μl ice-cold PBS. Protein concentration was measured using the BCA assay, adjusted to 2 mg/mL in ice-cold PBS (pH 7.4) and 50 μl was labelled with 2 µM Azido-fluorophosphonate (ActivX™ Azido-FP Serine Hydrolase Probe, Thermo Scientific) for 1 hour at room temperature. Fluorophosphonate-labelled proteins were conjugated to an alkyne IR800 dye (LI-COR, Inc) by click chemistry as described above. Samples were separated by SDS-PAGE, transferred to nitrocellulose for immunoblotting analysis, and visualised using a LI-COR Odyssey Imaging System.

### SDS-PAGE and Immunoblotting

All samples were heated for 5 minutes at 95 °C and vortexed before loading and resolving on 12% Tris-glycine SDS-PAGE gels. Proteins were detected by immunoblotting and fluorescent imaging using a LI-COR Odyssey M (LI-COR Biosciences) and quantified with Image Studio (LI-COR Biosciences).

### Computational Modelling of ABHD13 S193 protonation state

The AlphaFold2 (AF2) model of ABHD13 was downloaded from the AF Protein Structure database (https://alphafold.ebi.ac.uk/entry/Q7L211, accessed on 5 September 2023). After removing residues 1-42, part of the membrane-binding domain protruding from the αβ domain, the remaining structure (residues 43-337) was parameterised with the CHARMM36 force field^64^. The resulting system was prepared for simulations using in-house scripts based on HTMD2.3.2 ^65^ and VMD1.9.4^66^ frameworks. This multistep procedure performs the preliminary hydrogen atoms addition employing the pdb2pqr^67^ and PROPKA3^68^ software combination through the systemPrepare HTMD implementation^69^, considering a simulated pH of 7.4. All the histidine residues apart from H129 (predicted as ε tautomer) and H298 (predicted as protonated) were predicted in the tautomeric state δ. TIP3P water molecules^70^ were added to the simulation box (final dimensions 103 Å x 173 Å x 182 Å) using the VMD Solvate plugin 1.5 (VMD Solvate plugin, Version 1.5; http://www.ks.uiuc.edu/Research/vmd/plugins/solvate/). Finally, the overall charge neutrality was reached by adding Na^+^/Cl^−^ counter ions (final ionic concentration of 0.150 M) using the VMD Autoionize plugin 1.3 (Autoionize Plugin, Version 1.3; http://www.ks.uiuc.edu/Research/vmd/plugins/autoionize/). An alternative ABHD13 system where the catalytic S193 was modelled as deprotonated (net charge –1, CHARMM36 patch SERD) was prepared in the same way.

For both ABHD13 systems, the equilibration and production simulations were computed using the ACEMD3^71^ MD engine. Systems were equilibrated in isothermal-isobaric conditions (NPT) using the Monte Carlo barostat^72^ with a target pressure of 1 atm, the Langevin thermostat^73^ with a target temperature of 310 K, along with a low damping factor of 1 ps^−1^ and an integration time step of 2 fs. Clashes between atoms were reduced through 500 conjugate-gradient minimisation steps, followed by a 4 ns long MD simulation with a linearly-released positional constraint of 1 kcal mol^−1^ Å^−2^ on protein Cα atoms. Subsequently, 350 ns of productive MD simulation were performed with an integration time step of 4 fs in the canonical ensemble (NVT). The temperature was set at 310 K, using a thermostat damping of 0.1 ps^-1^ and the M-SHAKE algorithm^74^ to constrain the bond lengths involving hydrogen atoms. The cut-off distance for electrostatic interactions was set at 9 Å, with a switching function applied beyond 7.5 Å. Long-range Coulomb interactions were handled using the particle mesh Ewald summation method (PME)^75^ by setting the mesh spacing to 1.0 Å. Frames were saved every 100 ps.

RMSD of the catalytic triad side chains and the distance between the S193 side chain oxygen atom and H298 ε nitrogen atom indicated higher stability of the deprotonated S193 (Supplementary Fig. 9). Hence, this form of ABHD13 was modelled in the successive phases of work.

### Dynamic docking of SNAP25b (simplified system) to ABHD13

The SNAP25b full sequence AF2 model was downloaded from Uniprot (P60880, https://www.uniprot.org/uniprotkb/P60880) since this is the most complete structure available, and because the available X-ray and NMR structures (PDB codes: 1kil, 3rk2, 3rk3, 3r10 and 2nlt) align well with the AF2 model. Only residues 57-109, containing the MTD, were retained, and C85, 88, 90, and 92 were palmitoylated. The resulting SNAP25b structure (without any membrane model) was placed ∼40 Å away from the ABHD13 catalytic site (deprotonated S193 Fig. 5 c.). The resulting system was solvated, neutralised, and equilibrated as reported above (final box dimensions 151 Å x 90 Å x 111 Å). SNAP25b with the F84C mutation and palmitoylated C84 was prepared following the same procedure.

Multiple walker supervised molecular dynamics (mwSuMD)^75^ was employed to simulate molecular binding events between SNAP25b and ABHD13. mwSuMD is an adaptive sampling method for speeding up the simulation of binding events between molecules without introducing an energetic bias. Briefly, during mwSuMD, a series of batches (walkers) of short unbiased MD simulations are performed in parallel. After each batch is concluded, scores are computed based on user-defined metrics of the simulated system (e.g. distance, RMSD, number of contacts). The best short simulation in terms of score is selected and extended by seeding the same number of walkers, with the same duration as in the preceding step. For the dynamic docking between SNAP25b and ABHD13, three 500 ps walkers were seeded at every cycle. The distance between the carbonyl carbon atom of the palmitoyl group and the S193 side chain oxoanionic atom was supervised (SMscore) until reaching the value of less than 10 Å in 10 independent replicas for each modified Cysteine (40 replicas for the WT + 10 replicas for F84C, in total). After that, mwSuMD trajectories were extended by 100 ns using classic (unsupervised) MD to relax the system. Simulations where the distance between the carbonyl carbon atom of the palmitoyl group and the S193 side chain oxoanionic atom remained below 10 Å were considered productive.

### Complete SNAP25b – ABHD13 Systems Modelling

In constructing the complete ABHD13-SNAP25b system, the same criteria and methods as previously described were applied, except for utilising the AlphaFold2 models for both SNAP25B (https://www.uniprot.org/uniprotkb/P60880/entry, accessed on 1 July 2025) and ABHD13 (https://www.uniprot.org/uniprotkb/Q7L211/entry, accessed on 1 July 2025) embedded within a membrane and the type of ions employed.

To generate the complex and orient the proteins, both ABHD13 and SNAP25b structures were initially simulated alone, in two separate systems, in the presence of a membrane for 1 μs each using the previously defined settings. For composing the SNAP25b-ABHD13 system, and the separate systems, the positions of both structures relative to the membrane were subsequently obtained using the Positioning of Protein in Membrane server 2.0 ^45^. The most representative frames from both simulations were clustered using an in-house script employing a K-means clustering algorithm with the elbow method to identify the most representative conformations.

The membrane was constructed based on synaptic membrane models^76^, maintaining a composition of 40% phosphatidylcholine, 32% phosphatidylethanolamine, 12% phosphatidylserine, 5% phosphatidylinositol, and 10% cholesterol. The bilayer was modelled using the “Lipid Bilayer” service of CHARMM-GUI with a minimum surface area of 152 Å × 54 Å × 153 Å. The number of lipids and compositional percentages are reported in Table 1:

**Table 1.** Membrane composition for the SNAP25b-ABHD13 system simulated with mwSuMD.

| Residue | Total # | Top Leaflet # | Bottom Leaflet # | Top % | Bottom % | Average % | Difference from Experimental % | # Atoms |
| --- | --- | --- | --- | --- | --- | --- | --- | --- |
| CHL1 | 40 | 19 | 21 | 6.05 | 6.66 | 6.3 | -3.7 | 2960 |
| POPI | 25 | 15 | 10 | 4.77 | 3.17 | 3.97 | -1.03 | 3425 |
| POPE | 231 | 119 | 112 | 37.89 | 35.55 | 36.72 | 4.72 | 28875 |
| POPS | 84 | 47 | 37 | 14.96 | 11.74 | 13.35 | 1.35 | 10668 |
| POPC | 249 | 114 | 135 | 39.58 | 36.3 | 37.94 | -2.06 | 33366 |
| TIP3 | 9261 | 59628 | 12451 | 82.7 | 17.3 | N/A | N/A | 216237 |

The system charge was neutralised using Ca²⁺ and Cl⁻ ions to simulate the cellular environment in SNARE complexes^77^. The construction of the ABHD13-SNAP25b system was accomplished by positioning the two structures with their centres of mass at 65 Å to minimise initial steric clashes while maintaining a distance between the palmitoylated cysteines and the ABHD13 catalytic site below 30 Å.

The successive MD simulations were performed using OPENMM8.1.1^78^. During the equilibration, the L-BFGS (Limited-memory Broyden–Fletcher–Goldfarb–Shanno)^79^ Energy minimisation was performed in three stages with progressively stricter tolerances (10, 1, and 0.1 kJ/mol/nm). A thermalisation phase then gradually heated the system from 210 K to 310 K in 5 K increments (5,000 steps each) while all protein and membrane atoms were harmonically restrained (k = 50 kcal/mol/nm²). During the equilibration phase, positional restraints are gradually eased from 50 to 0 kcal/mol/nm² in increments of 0.2 kcal/mol/nm² over 120 ns, followed by 30 ns NPT, with a Monte Carlo barostat set at 1 bar^80^ using the Langevin Middle integrator^81^ with a 2 fs timestep and a friction coefficient of 0.002 ps⁻¹ at 310 K.

Three mwSuMD simulations were performed, monitoring the distance between the carbonyl carbon atom of Cys85 and the Ser193 side chain oxoanionic atom during time windows of 240 ps duration with a total of 8 walkers per cycle. After visual inspection, a final unsupervised simulation time of 200 ns was seeded from the last frame of replicas 1 and 2, which reached a distance lower than 5 Å, reassigning the atomic velocities.

### MD Analysis

Interatomic distances and root mean square deviations (RMSD) were computed with VMD1.9.4. Interatomic contacts and hydrogen bonds were detected using the GetContacts scripts tool (https://getcontacts.github.io), setting a hydrogen bond donor-acceptor distance of 3.3 Å and an angle value of 120° as geometrical cut-offs. Contacts and hydrogen bond persistency are quantified as the percentage of frames (over all the frames obtained by merging the different replicas) in which protein residues formed contacts or hydrogen bonds with the ligand. Volumetric maps of palmitoyl aliphatic chains were performed with VMD1.9.4. Distances between residues were computed using MDTraj^82^.

### Statistics and reproducibility

Statistical analyses were carried out using GraphPad Prism 10. An unpaired two-tailed Student’s *t*-test was used for direct comparison between two groups, whereas one-way ANOVA with Tukey’s or Dunnett’s post hoc tests, as appropriate, was used for comparisons among three or more groups. Data distributions were assessed for approximate normality prior to parametric testing. Data are represented as means ± standard error of the mean (SEM).

## DATA AVAILABILITY

All data reported in this paper will be shared by the lead contact upon request. mwSuMD trajectories of the full-length SNAP25b:ABHD13 complex (Replicas 1 and 2) are available at Zenodo’s webpage https://zenodo.org/records/18233546

---

**Supplementary Fig. 1:**
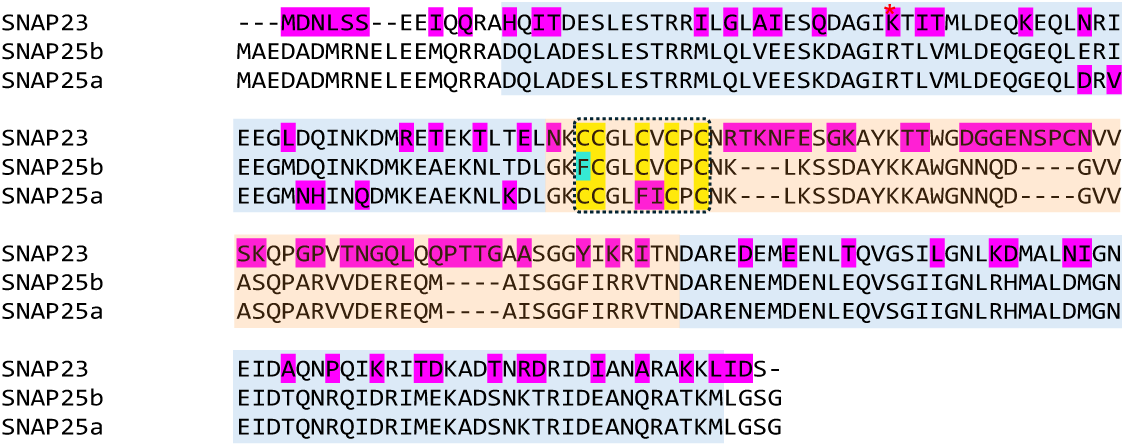
Sequence alignment of SNAP25 isoforms. Protein sequence alignment of human SNAP25b (P60880-1), SNAP25a (P60880-2) and SNAP23 (O00161). The two SNARE domains are shaded in blue. The linker domain is shaded in peach with *S*-acylated cysteines highlighted in yellow. The amino acids shaded in magenta in SNAP25a and SNAP23 differ from SNAP25b. A single phenylalaline shaded in turquoise (F84) is the only deviation in SNAP25b not shared with SNAP25a or SNAP23. The protein sequence alignments were generated using Clustal Omega^76^.

**Supplementary Fig. 2:**
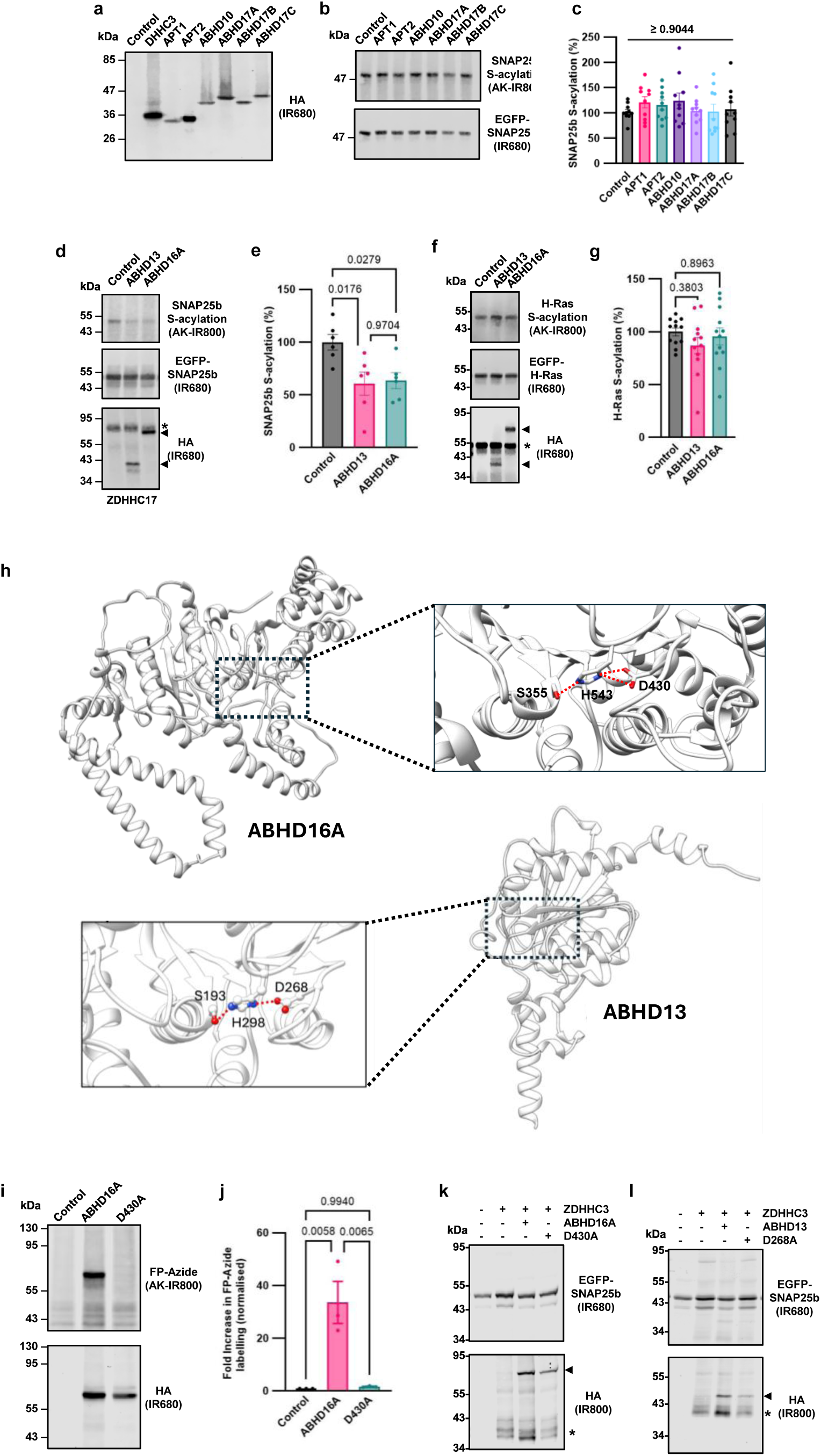
SNAP25b deacylation. **a.** Expression of HA-ZDHHC3 and HA-tagged metabolic serine hydrolase thioesterases in transfected HEK293T cells. **b.** Representative immunoblot images of SNAP25b *S*-acylation levels following co-expression of HA-ZDHHC3 and HA-tagged APT1, APT2, ABHD10, ABHD17A, ABHD17B, ABHD17C, or empty vector control. **c.** Quantification of SNAP25b *S*-acylation normalized to control (ZDHHC3 and empty vector). **d.** Representative immunoblot images of SNAP25b *S*-acylation levels following co-expression of HA-ZDHHC17 and HA-tagged ABHD13, ABHD16A, or empty vector control. **e.** Quantification of SNAP25b *S*-acylation normalized to control (ZDHHC17 and empty vector). **f.** Representative immunoblot images of H-Ras *S*-acylation levels following co-expression of HA-tagged ABHD13, ABHD16A, or empty vector control. **g.** Quantification of H-ras *S*-acylation normalized to empty vector control. **h.** AlphaFold structural models of human ABHD16A and ABHD13 highlighting the predicted the catalytic triad. **i.** HEK293T total membrane proteomes expressing HA-tagged ABHD16A, ABHD16A(D430), or empty vector (control) were labelled with ActivX Azido-FP Serine Hydrolase Probe and conjugated to an alkyne-infrared 800 dye (AK-IR800) by click chemistry. Representative immunoblot images of Azido-FP labelling. **j.** Quantification of Azido-FP binding represented as a fold change compared with empty vector. **k.** Representative immunoblot images of EGFP-SNAP25b, HA-ZDHHC3 and HA-tagged ABHD16A WT and D430A expression levels prior to membrane fractionation. **l.** Representative immunoblot images of EGFP-SNAP25b, HA-ZDHHC3 and HA-tagged ABHD13 WT and D268A expression levels prior to membrane fractionation. All *p* values obtained by one-way ANOVA Tukey’s multiple comparison test.

**Supplementary Fig. 3:**
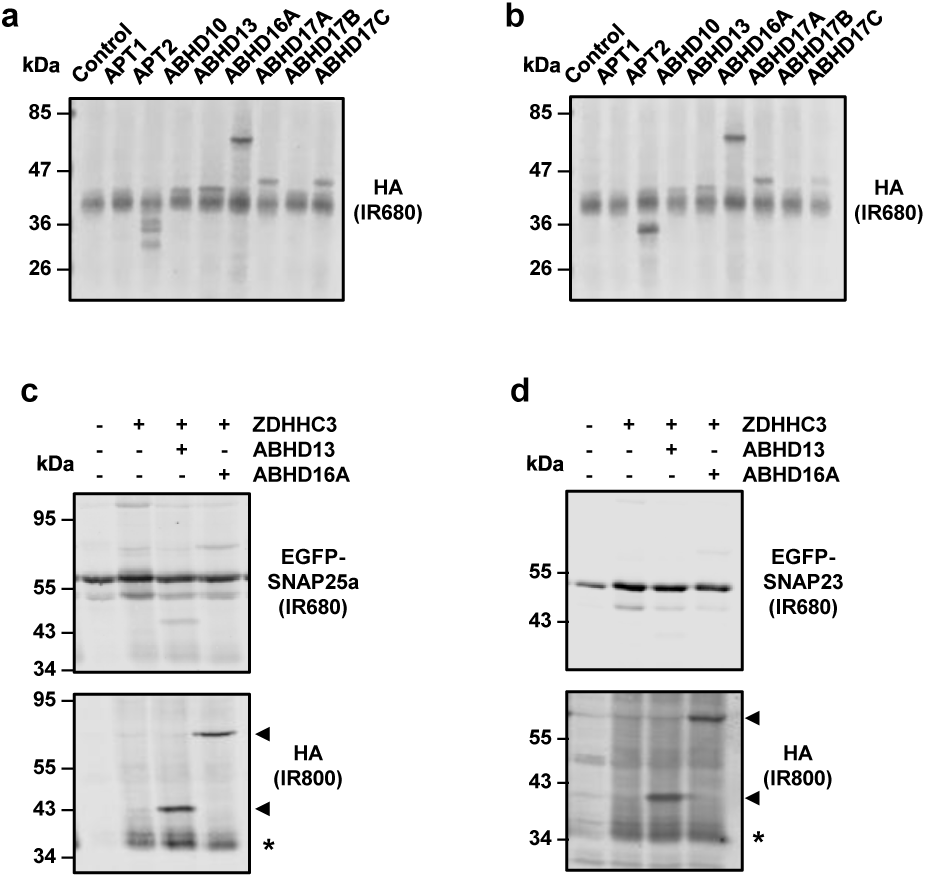
SNAP25a and SNAP23 deacylation and membrane fractionation. **a, b.** Representative immunoblot images of co-expressed HA-tagged deacylases and ZDHHC3 for SNAP25a (a) or SNAP23 (b) shown in Figure 4 a and b. **e. f.** Representative immunoblot images of EGFP-SNAP25a (c.) or EGFP-SNAP23 (d.), HA-ZDHHC3 and HA-tagged ABHD13 or ABHD16A expression levels prior to membrane fractionation shown in Figure 4 e and f.

**Supplementary Fig. 4:**
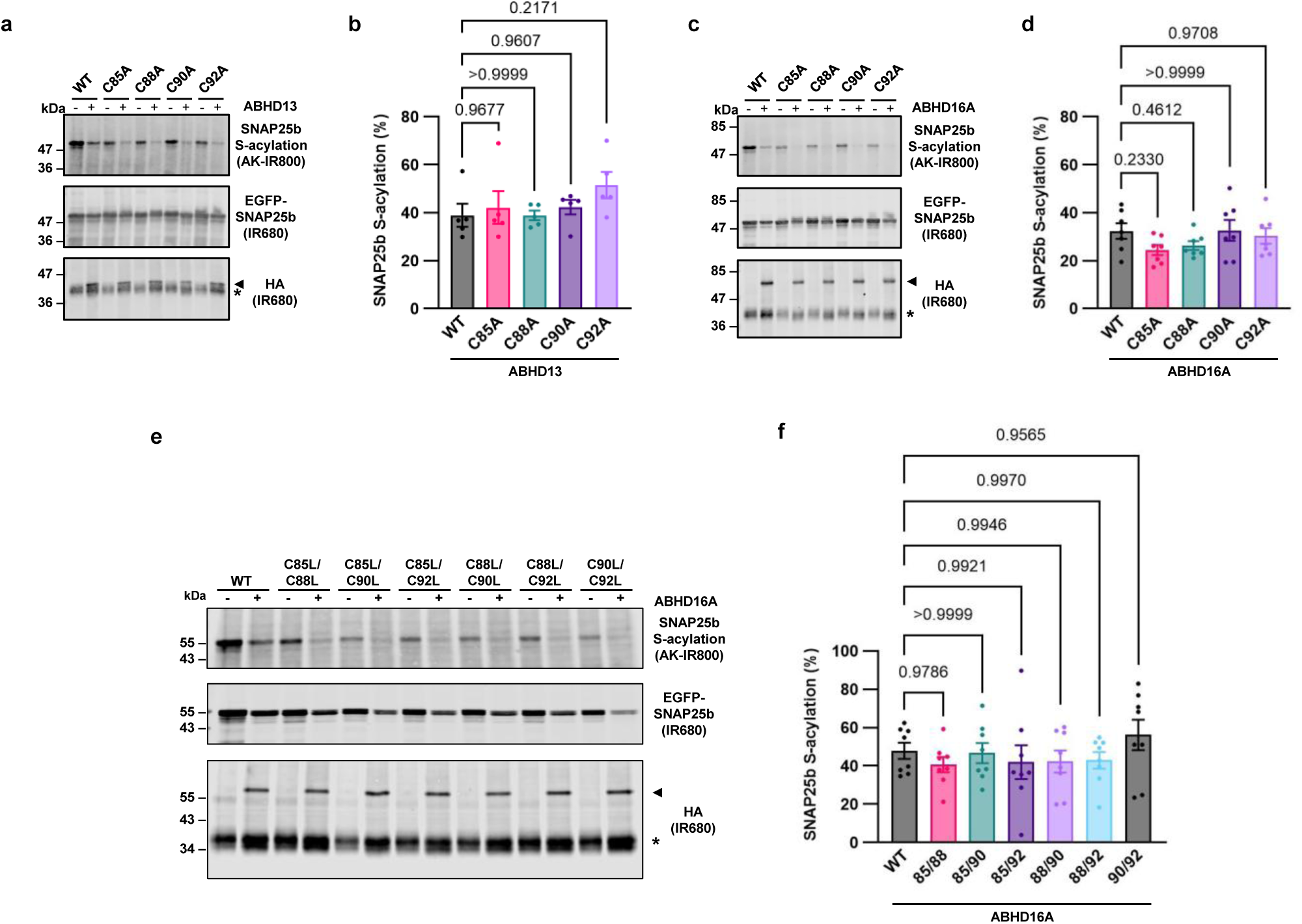
Deacylation of SNAP25b cysteine mutants by ABHD16A and ABHD13. **a., c, e.** *S*-acylation levels of SNAP25b WT, single (**a.** and **c.**) or double cysteine substitutions (**e.**) co-transfected with ZDHHC3 and empty vector (control), ABHD16A (**a., c, e.)** or ABHD13 (**a. and c.)** determined by C16:0-azide-labelling, click chemistry and immunoblotting**. b., d., f.** Quantification of SNAP25b *S*-acylation levels shown in a., c., e.

**Supplementary Fig. 5:**
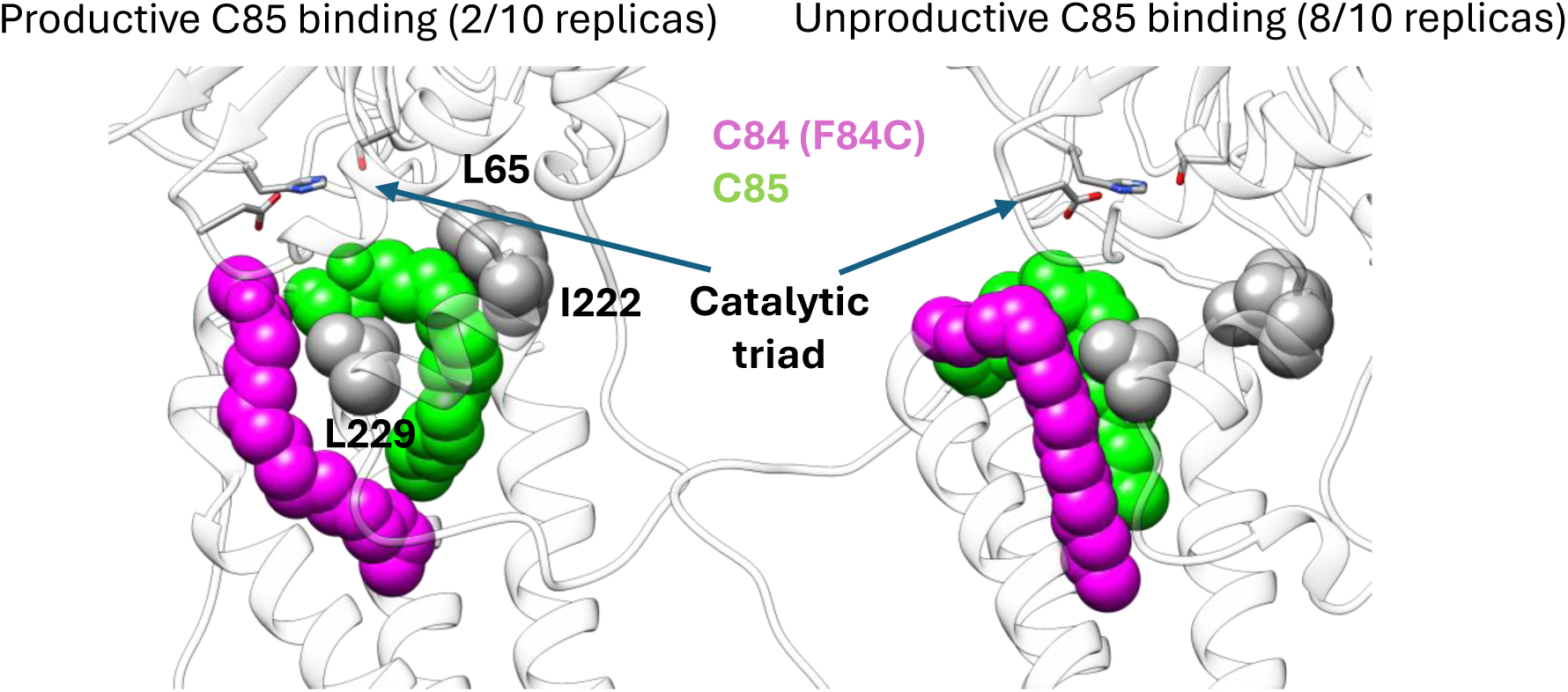
The F84C palmitoyl group steric hindrance disfavours C85 recognition during mwSuMD (simplified system).

**Supplementary Fig. 6:**
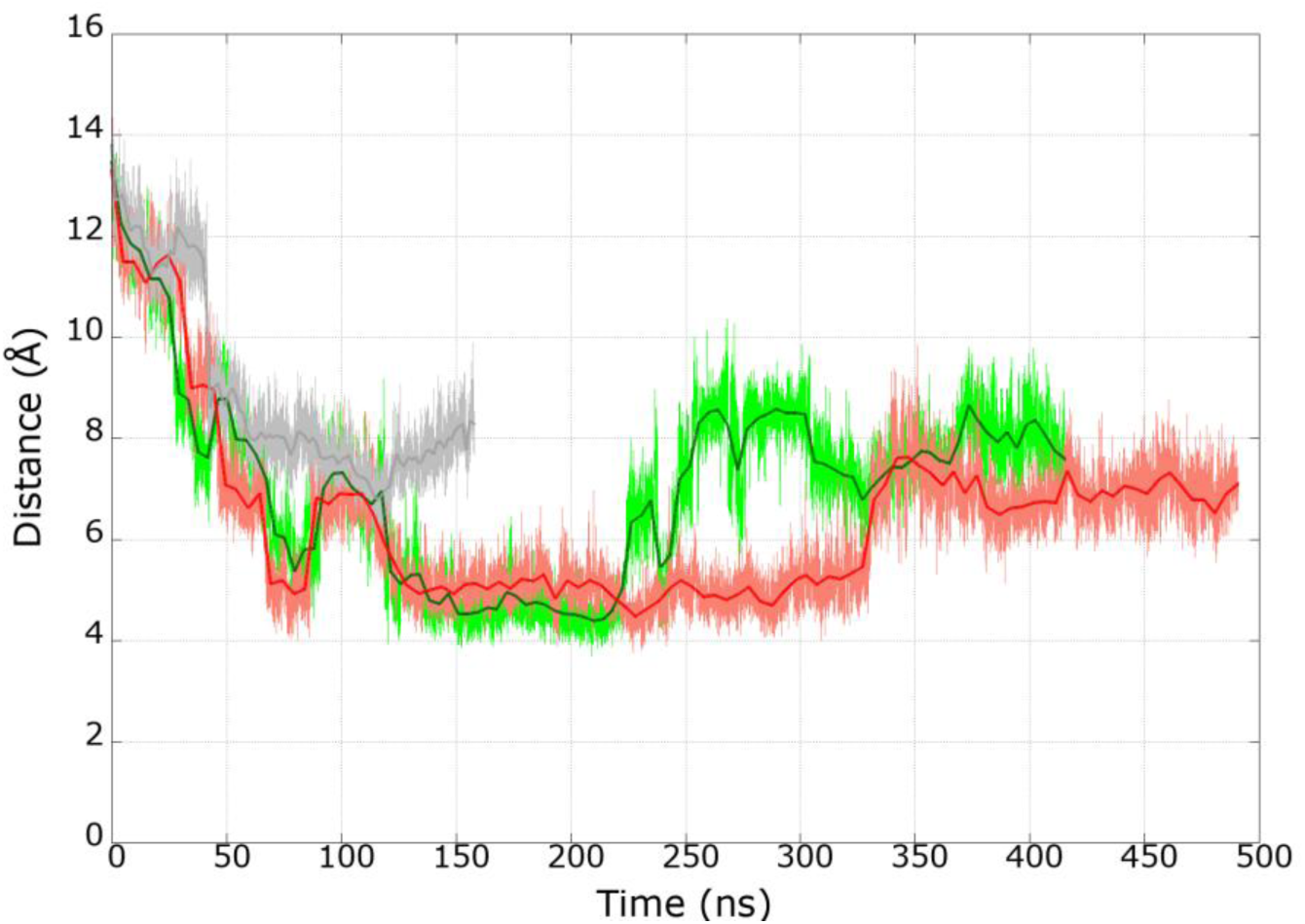
Distance between the carbonyl carbon atom of C85 and the S193 oxygen atom during mwSuMD simulations of the complete system.

**Supplementary Fig. 7:**
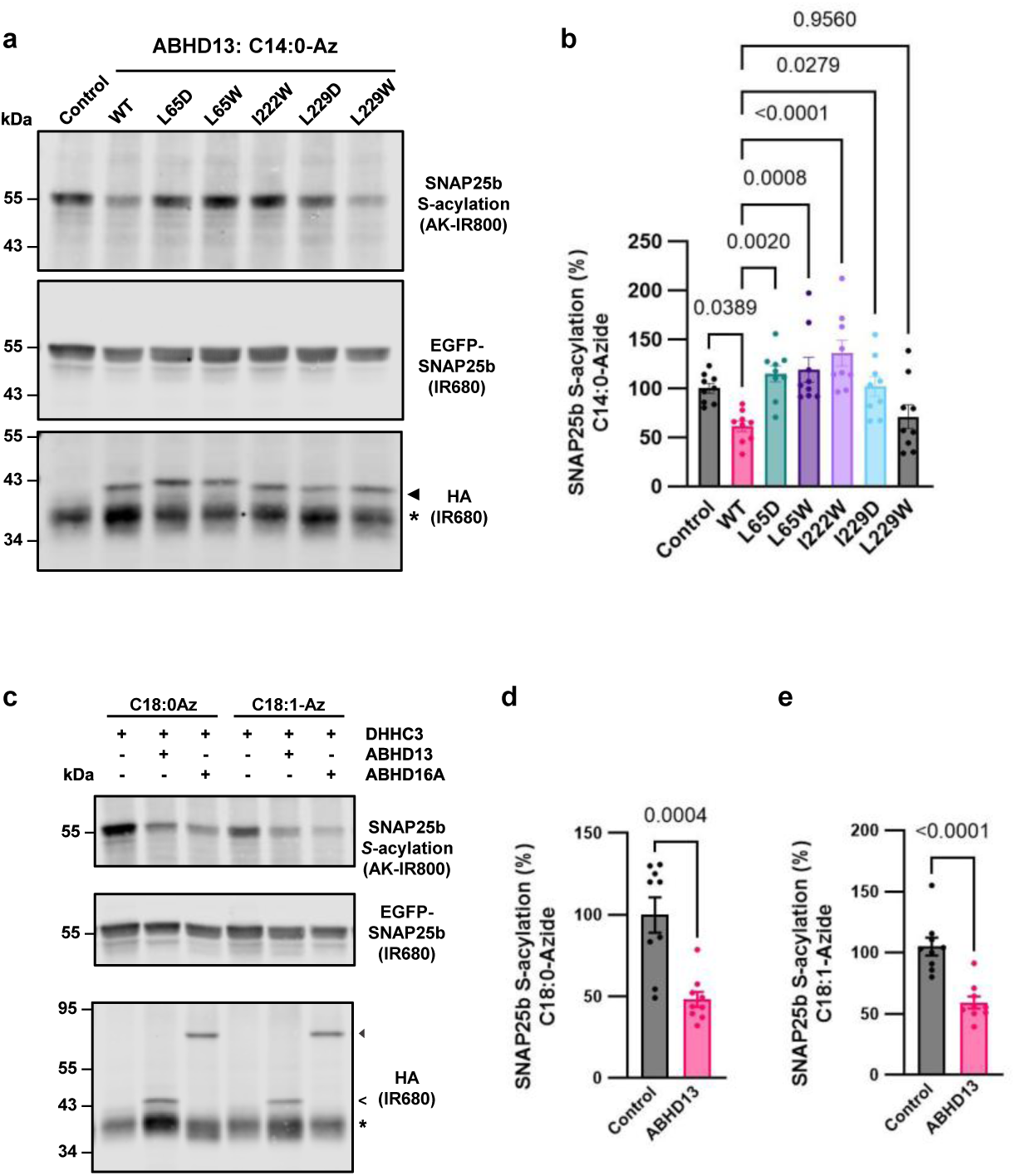
ABHD13-mediated deacylation of SNAP25b with various fatty acid-azide substrates. **a.** S-acylation levels of SNAP25b co-transfected with ZDHHC3 and empty vector (control), ABHD13 WT or pocket mutants determined by C14:0-azide-labelling, click chemistry and immunoblotting. **b.** Quantification of SNAP25b S-acylation levels shown in **a. c.** S-acylation levels of SNAP25b co-transfected with ZDHHC3 and empty vector (control) or ABHD13 determined by C18:0 or C18:1-azide-labelling, click chemistry and immunoblotting. **d. e.** Quantification of SNAP25b S-acylation levels shown in **c.**

**Supplementary Fig. 8:**
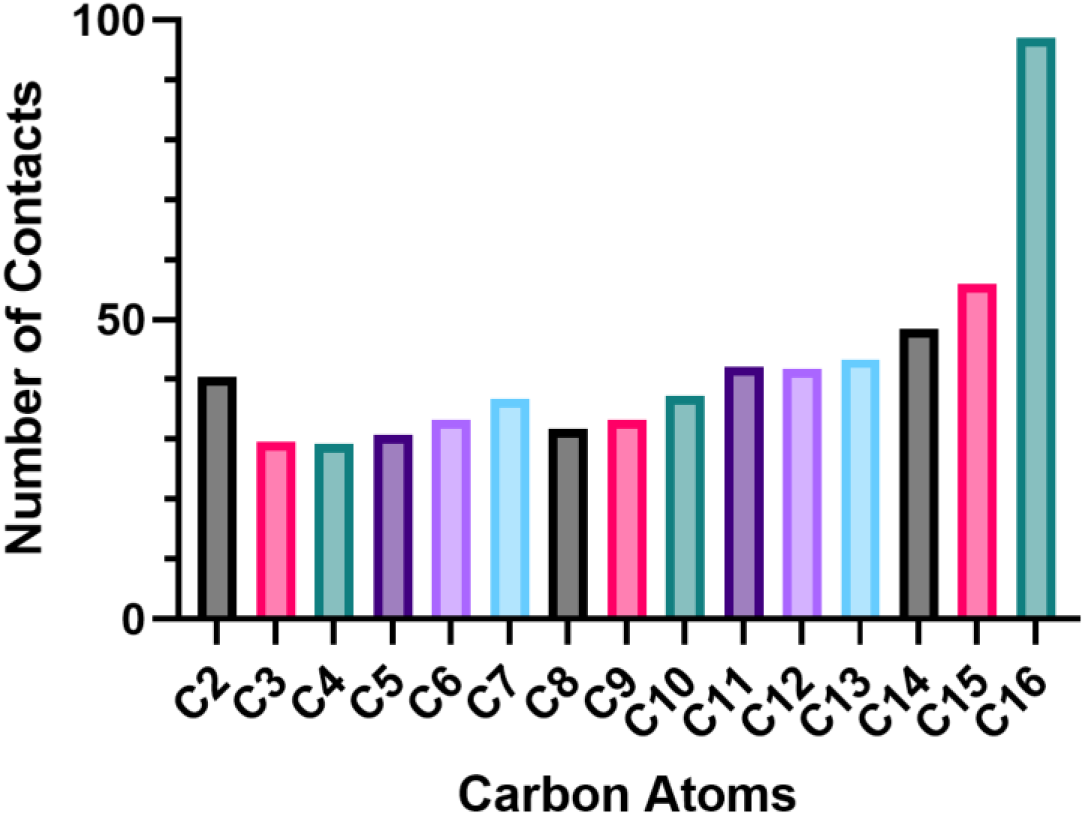
Hydrophobic atomic contacts during mwSuMD of full-length ABHD13 and SNAP25b. Contacts are expressed as the sum of contacts for each carbon atom, divided by the total number of frames, multiplied by 100.

**Supplementary Fig. 9:**
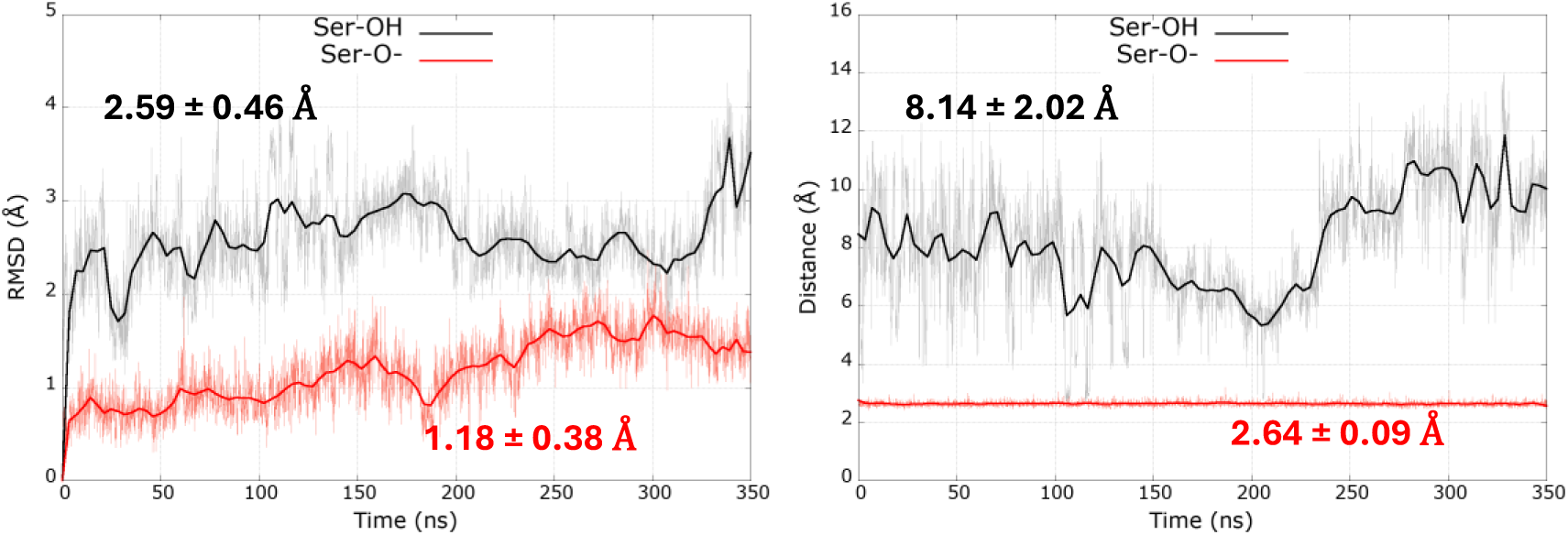
Comparison between the deprotonated (red) and the protonated Ser193 (black) during explorative classic MD. Left-hand panel: RMSD of the ABHD13 cytosolic domain; right-hand panel: distance between Ser193 side chain oxygen atom and His298 epsilon nitrogen atoms.

## Supplementary Videos

**Video S1.** Dynamic docking between SNAP25b (magenta) and ABHD13 (white). Palmitoyl groups are shown as sticks, while ABHD13 catalytic residues and side chain interacting with the C85 palmitoyl group are shown as cyan sticks.

**Video S2.** mwSuMD simulations (replica 2) showing the recognition of SNAP25b C85 palmitoyl group (red van der Waals representation) by ABHD13 hydrophobic pocket residues L65, I22, and L229 (thick white ribbon). Membrane polar heads’ nitrogen atoms are displayed as green spheres; S193, H298, and D268 are shown as thin white ribbons for reference; C88, C90, and C92 palmitoyl groups are in a thin, red stick representation.

